# Natural Variation in Maize Shikimate Dehydrogenase Alters Enzyme Activity and Kernel Homoserine Accumulation

**DOI:** 10.64898/2026.08.26.747334

**Authors:** Rajnee Hasan, Gen Xu, Taiwo Dele-Osibanjo, Niaz Bahar Chowdhury, Connor Pedersen, Rajib Saha, Jinliang Yang, Toshihiro Obata

**Author notes:** Address correspondence to: Toshihiro Obata, University of Nebraska-Lincoln, Department of Biochemistry and Center for Plant Science Innovation 1901 Vine Street, Lincoln, NE, 68588.

## Abstract

Metabolic diversity in maize kernels determines nutritional quality and end-use value. Therefore, understanding its genetic basis is essential for crop improvement and elucidating plant metabolic regulation. Here, we integrated metabolite profiling with metabolite-based genome-wide association studies (mGWAS), structural modeling, enzyme kinetics, and genome-scale metabolic simulations to identify genetic determinants of kernel metabolite variation in 265 maize inbred lines. Profiling of 57 metabolites revealed inter-genotypic variation, with homoserine among the most variable metabolites. mGWAS identified 62 locus–trait associations implicating 788 candidate genes, including 154 encoding metabolic enzymes. A major association for homoserine mapped to the shikimate dehydrogenase gene *Sad1* on chromosome 10. Four tightly linked coding-region SNPs, including three non-synonymous variants, defined two *Sad1* haplotypes associated with differential homoserine accumulation, independent of gene expression variation. Structural analysis and recombinant enzyme assays showed that these substitutions occur within catalytic and cofactor-binding domains and alter catalytic efficiency. Genome-scale metabolic modeling indicated that variation in SAD1 activity influences plastidial oxaloacetate availability for aspartate and homoserine biosynthesis through redox-coupled flux via the malate-oxaloacetate shuttle. Together, our results indicate that *Sad1* allelic variation alters enzyme function and amino acid accumulation, linking the shikimate pathway, redox metabolism, and amino acid biosynthesis in maize kernels.

## Introduction

Maize (*Zea mays* L.) is a globally significant crop, serving as a primary source of calories, animal feed, and industrial raw materials. The nutritional and industrial value of maize grain is influenced by the metabolic composition of its kernels, which contain diverse primary and specialized metabolites involved in plant growth, development, defense, and grain quality. Previous studies have demonstrated substantial metabolic diversity in maize cultivars, including differential accumulation of flavonoids in leaves (Zhou et al. 2019) and kernels (Wen et al. 2014), expanded these investigations to additional germplasm panels, including edible maize and globally diverse kernel populations, to identify numerous genetic loci associated with metabolite variation. However, the genetic factors underlying natural variation in kernel metabolites remain incompletely understood. Identifying these factors is important for improving maize nutritional quality, stress resilience, and end-use value through targeted breeding.

Metabolite-based genome-wide association studies (mGWAS) provide a powerful approach for dissecting the genetic architecture of metabolic traits. By linking metabolite abundance with genome-wide polymorphisms, mGWAS can identify metabolite quantitative trait loci (mQTLs), which are genomic regions associated with variations in metabolite accumulation. mGWAS has been successfully applied in maize (Riedelsheimer et al. 2012), rice (Wang et al. 2018), wheat (Chen et al. 2020), soybean (Zhou et al. 2015), sorghum (Zheng et al. 2011) and cotton (Du et al. 2018), revealing genetic determinants of metabolite accumulation and pathway regulation. In the present study, we used maize kernels from 265 inbred lines from the Maize Association Panel (MAP), which was designed to represent up to 80% of the genetic diversity found within maize (Flint-Garcia et al. 2005; Mural et al. 2022) and has been shown to exhibit substantial chemical diversity (Zhou et al. 2019).

One of the key strengths of mGWAS is its ability to identify candidate metabolic enzyme-coding genes underlying natural variation in metabolite abundance. Variants within these candidate genes provide opportunities for functional characterization, enabling possibly mechanistic studies of how genetic variation alters enzyme activity. For instance, a polymorphism in exon 7 of the *BADH2* gene produces a nonfunctional enzyme, leading to an accumulation of the aroma precursor 2-acetyl-1-pyrroline in rice (Giang et al. 2023). In maize, polymorphisms in a tocopherol methyltransferase gene are associated with variation in α-tocopherol levels (Li et al. 2012). Coding-sequence polymorphisms can cause amino acid substitutions in enzyme active sites and structural domains, thereby affecting enzyme activities by altering enzyme abundance, stability, substrate affinity, cofactor preference, or catalytic efficiency. For example, substitution of tyrosine with phenylalanine in the active site of cyclodextrin glucanotransferases in *Bacillus ohbensis* changed its activity and increased cyclodextrin production (Leemhuis et al. 2010). Thus, coding variations in metabolic enzymes can provide a direct mechanistic link between genetic polymorphisms and metabolic phenotypes.

The shikimate pathway represents a critical metabolic junction in plants, fungi, and bacteria, bridging primary carbon metabolism and aromatic amino acid and specialized metabolite biosynthesis (Pott et al. 2019). This seven-step enzymatic cascade begins with phosphoenolpyruvate from glycolysis and erythrose 4-phosphate from the pentose phosphate pathway, thereby funneling carbon from core energy and biosynthesis pathways. Shikimate dehydrogenase catalyzes the fourth step of the pathway, the NADPH-dependent reduction of 3-dehydroshikimate to shikimate (Herrmann 1995). The pathway ultimately produces chorismic acid, a branch-point intermediate for aromatic amino acids-phenylalanine, tyrosine, and tryptophan, and a vast array of aromatic specialized metabolites involved in plant defense, structural integrity, and signaling (Shende et al. 2024). Since the shikimate pathway competes for central carbon precursors (Maeda and Dudareva 2012; Yokoyama et al. 2021), natural variation in its enzymes may influence not only aromatic amino acid metabolism but also the allocation of precursors to other metabolic routes.

While mGWAS identifies numerous statistical associations between genomic loci and metabolic phenotypes, establishing the underlying causal mechanisms through functional interpretation and experimental validation remains the primary challenge. Metabolic network modeling provides a complementary systems-level approach by mathematically representing known biochemical reactions as an integrated network that can connect genes, enzymes, metabolites, and metabolic fluxes (Clark et al. 2020). These models can predict how changes in enzyme activity or pathway capacity affect metabolic flux distribution and metabolite accumulation (Yuan et al. 2016). Metabolic network models have been applied to various plant studies, including abiotic stress responses in rice (Lakshmanan et al. 2013, 2015; Poolman et al. 2013) and C4 metabolism in maize (Saha et al. 2011; Gomes De Oliveira Dal’Molin et al. 2018), storage metabolism in barley seeds (Grafahrend-Belau et al. 2009), and monoterpene biosynthesis in glandular trichomes (Johnson et al. 2017). In this study, we used a maize kernel metabolic network model (Chowdhury et al. 2023) to evaluate the metabolic link between an mGWAS-associated enzyme-coding gene and metabolic phenotype.

In this study, we combined GC-MS primary metabolite profiling, mGWAS, structural modeling, enzyme kinetic analysis, and metabolic network modeling to dissect the genetic basis of the maize kernel metabolic variation in the maize diversity panel. Analysis of 57 metabolites identified multiple metabolite-associated loci and highlighted a significant association between kernel homoserine accumulation and natural variations in the *Shikimate dehydrogenase 1* (*Sad1*) gene. Enzyme kinetics and functional modeling analyses suggest that *Sad1* allelic variation alters enzyme activity and affects plastidial redox metabolism, affecting the metabolic flux to aspartate-family amino acid biosynthesis. These findings provide a mechanistic framework linking natural genomic variation to kernel metabolite accumulation in maize.

## Materials and Methods

### Field experimental design and sample collection

A subset of 265 genotypes from the maize diversity panel (Flint-Garcia et al. 2005; Table S1) was analyzed in this study. This panel represents a broad sampling of the genetic diversity present in cultivated maize, providing a robust foundation for exploring natural variation in seed metabolite composition (Harrigan et al. 2007; Skogerson et al. 2010). Plants were grown under rainfed conditions in a field previously planted with commercial maize at the Havelock Research Farm of the University of Nebraska-Lincoln. The kernels were harvested during the 2019 growing season (Palali Delen et al. 2023). Urea (120 lbs acre^-1^) was applied as a nitrogen source before planting. Each genotype was planted in a two-row sub-plot measuring 20 ft × 5 ft (6.096 m × 1.524 m), with 30 inches (76.2 cm) of spacing between rows and 6 inches (15.24 cm) of spacing between plants within a row. Each row was sown with 38 seeds, corresponding to an approximate planting density of 30,000 seeds per acre. At maturity, kernels were harvested from all plants within each genotype’s sub-plot. For each genotype, one composite sample consisting of 10 kernels was randomly selected from the pooled harvest to serve as the representative biological sample for metabolite extraction.

### Sample preparation and metabolite profiling

Metabolites were extracted from maize kernel tissue using a biphasic methanol– chloroform–water protocol adapted from Wase et al. (Wase et al. 2022). Briefly, 50 mg of homogenized maize kernel powder was extracted with 700 µL of 99.9% (v/v) methanol containing ribitol (0.2 mg/mL) as an internal standard. Phase separation was induced using chloroform: water (2:1, v/v), and the upper aqueous phase was collected. Extracts were dried completely in a SpeedVac concentrator without heat to prevent degradation of thermolabile metabolites and stored at −80 °C prior to derivatization. For GC-MS analysis, dried extracts were derivatized by methoxyamination with methoxyamine hydrochloride in pyridine (20 mg/mL) at 37 °C for 2 h, followed by trimethylsilylation using MSTFA containing FAMEs (50:1) at 37 °C for 30 min. Derivatized samples were transferred to autosampler vials for analysis.

Metabolite profiling was performed using a 7200 GC-QTOF system (Agilent Technologies, Santa Clara, CA) equipped with a DB-5MS column (30 m × 0.25 mm × 0.25 μm) operating in both splitless and split mode. Helium was used as the carrier gas at a constant flow rate of 1 mL/min. The oven temperature was programmed from 80 °C to 330 °C at 15 °C/min and held at 330 °C for 6 min. Mass spectra were acquired in electron ionization mode at 70 eV over a mass range of 85–450 m/z. Raw data were processed using MassHunter software version B.08.00 (Agilent Technologies) for peak detection and deconvolution. Metabolites were annotated by comparison with the Fiehn library (Kind et al. 2009) based on mass spectra and retention indices using MassHunter Unknowns

Analysis software and quantified using and Quantitative Analysis software, followed by extensive manual curation. Relative metabolite abundances were calculated by normalizing peak ion intensities to the internal standard (ribitol) and the precise sample fresh weight.

### Metabolite-based genome-wide association mapping

We conducted GWAS for 57 metabolites using the GEMMA software (Zhou and Stephens 2012) with 22.5 million SNPs. In the analysis, we fitted the first 3 principal components (PCs) as covariates calculated from genome-wide SNPs using the PLINK software (Chang et al. 2015). To account for the genetic relatedness, we used the centered identical by state kinship matrix generated by TASSEL software (Bradbury et al. 2007) as the covariance matrix as random effects in the GWAS model. The threshold for the significant association SNPs was set to 6.5 ×10^-8^ (0.05/n, n = 769,690 is the number of independent SNPs). Here, the independent SNP number was determined by using PLINK 1.9 with the indep-pairwise option (window size 10 kb, step size 10, *r^2^* ≥ 0.1). GWAS peaks were then determined by considering a window 500 kb upstream and downstream of the significant SNPs. Overlapping regions were merged. For mGWAS data analysis, first, we selected associations from genomic locations with signal counts exceeding 5% of the maximum number of signals (N) detected at a single location. Second, we independently selected associations within most significant 5% based on P value. The overlap of these two filtered sets were chosen for locus-trait association analysis.

### Vector construction, protein expression and purification

The open reading frame sequence of *Z. mays Sad1* Hap1 was designed according to the *Sad1* sequence in B73 inbred after eliminating the genomic sequence encoding plastidial signal peptide. The Hap2 construct contained sequence variations causing specific amino acid substitutions resembling Hap2, namely F46L, P522S, and L537Q. Both constructs were designed to fuse an HA epitope tag to the C-terminus of both *Sad1* Hap1 and *Sad1* Hap2 for immunodetection. The DNA fragments were synthesized and cloned into the NdeI/XhoI sites of the pET21b expression vector by GenScript (Piscataway, NJ, USA) (Supplementary doc 1). The expression vectors, designated pET-Sad1_Hap1 and pET-Sad1_Hap2, were transformed into chemically competent *Escherichia coli* BL21 (DE3) cells for heterologous protein expression.

The resulting *E*. *coli* strains were cultured in Terrific Broth (TB) medium at 37°C with shaking at 220 rpm until the OD_600_ reached 0.6–0.8. Recombinant protein expression was induced by the addition of 0.5 mM isopropyl-β-D-thiogalactopyranoside (IPTG), followed by an additional 20 h of incubation at 18°C. Cells were harvested by centrifugation (6,000 x *g*, 20 min, 4 °C), and the resulting pellets were resuspended in 5 volumes (w/v) of ice-cold lysis buffer comprising 50 mM Tris-HCl (pH 8.0), 500 mM NaCl, 10% (v/v) glycerol, 20 mM imidazole, 200 µg mL^-1^ lysozyme, 20 µg mL^-1^ DNase I, 1 mM MgCl_2_, 0.5% (v/v) Tween 20, 10 mM dithiothreitol (DTT), and one tablet of EDTA-free protease inhibitor cocktail per 100 mL lysis buffer.

Cells were lysed by sonication using a Sonic Dismembrator 550 (Fisher Scientific, Hampton, NH, USA) at an amplitude setting of 6, operating with a 5 sec on / 15 sec off pulse cycle for a total process time of 7 min. Cellular debris was pelleted by centrifugation at 16,000 x g for 70 min at 4 °C. The target recombinant proteins in the supernatant were purified using Ni-NTA His SpinTrap columns (GE Healthcare, Chicago, IL, USA). The columns were washed sequentially with a step gradient of imidazole [40 mM, 60 mM, and 80 mM in a base purification buffer of 50 mM Tris-HCl (pH 8.0), 500 mM NaCl, and 10 mM β-mercaptoethanol], and the HA-tagged proteins were subsequently eluted using the same buffer supplemented with 500 mM imidazole (Figure S2). The eluted proteins were desalted six times with 10 mL of 50 mM Tris-HCl (pH 8.0) using an Amicon Ultra-4 centrifugal concentrator (10 kDa MWCO; MilliporeSigma) at 4 °C. Purified proteins were evaluated by SDS-PAGE with Coomassie Brilliant Blue staining, and western blotting using anti-HA tag monoclonal antibody (LSG26183, Thermo Fisher Scientific) as primary antibody (Figure S2). Purified SAD1 protein solutions showed multiple bands in both staining, indicating partial impurity of the proteins. Nevertheless, we used these samples for enzyme activity assays because the apparent protein band pattern in these samples were identical and unlikely affect the enzyme activity (Figure S2). Glycerol was appended to a final concentration of 10% (v/v) before the purified enzymes were aliquoted and flash-frozen in liquid nitrogen and stored at-80°C prior to downstream analyses. Total protein concentrations were quantified using the Bio-Rad Protein Assay Kit (Bio-Rad Laboratories, Hercules, CA, USA) according to the manufacturer’s instructions.

### Enzyme activity and kinetics assay

Shikimate dehydrogenase enzymatic activities of both SAD1-Hap1 and SAD1-Hap2 were determined by monitoring the formation of NADPH through its absorption at 340 nm (ε340 = 6.22 mM^-1^ cm^-1^), adapting previously established protocol (Zhang et al. 2005; Singh and Christendat 2006; Díaz-Quiroz et al. 2018; Almeida et al. 2024) with minor modifications. Assays were executed in a 200 µL total volume containing 100 mM Tris-HCl buffer (pH 9.0), 5 mM MgCl_2_, 2 mM NADP^+^, and 0.118 nM of purified enzyme. Following component assembly and thermal equilibration at 25°C, the reaction was initiated by the addition of 5 mM shikimic acid (Sigma-Aldrich, St. Louis, MO, USA). The initial linear rate of NADP^+^ reduction was tracked for 10 min using a CLARIOstar Plus microplate reader (BMG LABTECH, Ortenberg, Germany). One unit of SAD1 activity was defined as the quantity of enzyme required to generate 1 µmol of NADPH per minute. Specific catalytic activity was normalized against total protein content and expressed as U mg^-1^ (µmol min^-1^ mg^-1^ protein).

The determination of the Michaelis–Menten constant (*K*_m_), maximum velocity (*V*_max_), and turnover number (*k*_cat_) for both Hap1 and Hap2 was performed by varying the concentrations of NADP^+^ while keeping the shikimic acid concentration of 5 mM. The reaction was initiated by addition of 0.118 nM of enzyme. NADP^+^ concentrations were 0.05, 0.1, 0.15, 0.2, 0.25, 0.3, 0.5, 1, 2, 3, 4 and 5mM. The *K*_m_, *V*_max_, and *k*_cat_ were calculated from double reciprocal Lineweaver–Burk plots.

### Maize kernel metabolic model reconstruction and shikimate dehydrogenase flux simulation

Flux balance analysis (FBA) (Orth et al. 2010) and flux sum analysis (FSA) (Chowdhury et al. 2022) were performed using the maize seed metabolic model derived from the multi-organ maize metabolic model iZMA6517(Chowdhury et al. 2023) using the General Algebraic Modeling System (GAMS) version 24.7.4 with the IBM CPLEX solver. FBA was first used to maximize seed biomass production, after which biomass was constrained to 99% of its maximum value while the shikimate dehydrogenase fluxes were systematically varied to evaluate homoserine metabolite pool size and, therefore, their theoretical effects on the aspartate and shikimate branches. Finally, flux distribution was calculated using the highest possible flux of shikimate dehydrogenase.

## Results

### Kernels in the maize diversity panel exhibit a diverse metabolite profile

Using high-throughput GC-MS metabolite profiling, we quantified the relative levels of 57 metabolites in mature maize kernel extracts from the 265 inbred lines of the maize diversity panel (Table S1). To assess the metabolic diversity available for genetic mapping, we calculated the standard deviation (SD) of the relative abundance for each metabolite across the panel (Figure 1A). We observed a broad range of metabolic variability among the 57 metabolites, with SD values spanning from 0.29 to 2.57. The metabolites exhibiting the highest variation with SD greater than 1.5 included glyoxylic acid, cysteine, trehalose, dihydroxybenzoic acid, mannitol, galactinol, and homoserine. The corresponding abundance distributions showed that these high-SD metabolites displayed wide interquartile ranges, reflecting significant segregation of these metabolic phenotypes within the diversity panel (Figure 1B). In contrast, metabolites such as ferulic acid, palmitic acid, and stearic acid were highly conserved (SD < 0.5), showing narrow distributions clustered tightly around the mean.

**Figure 1.**
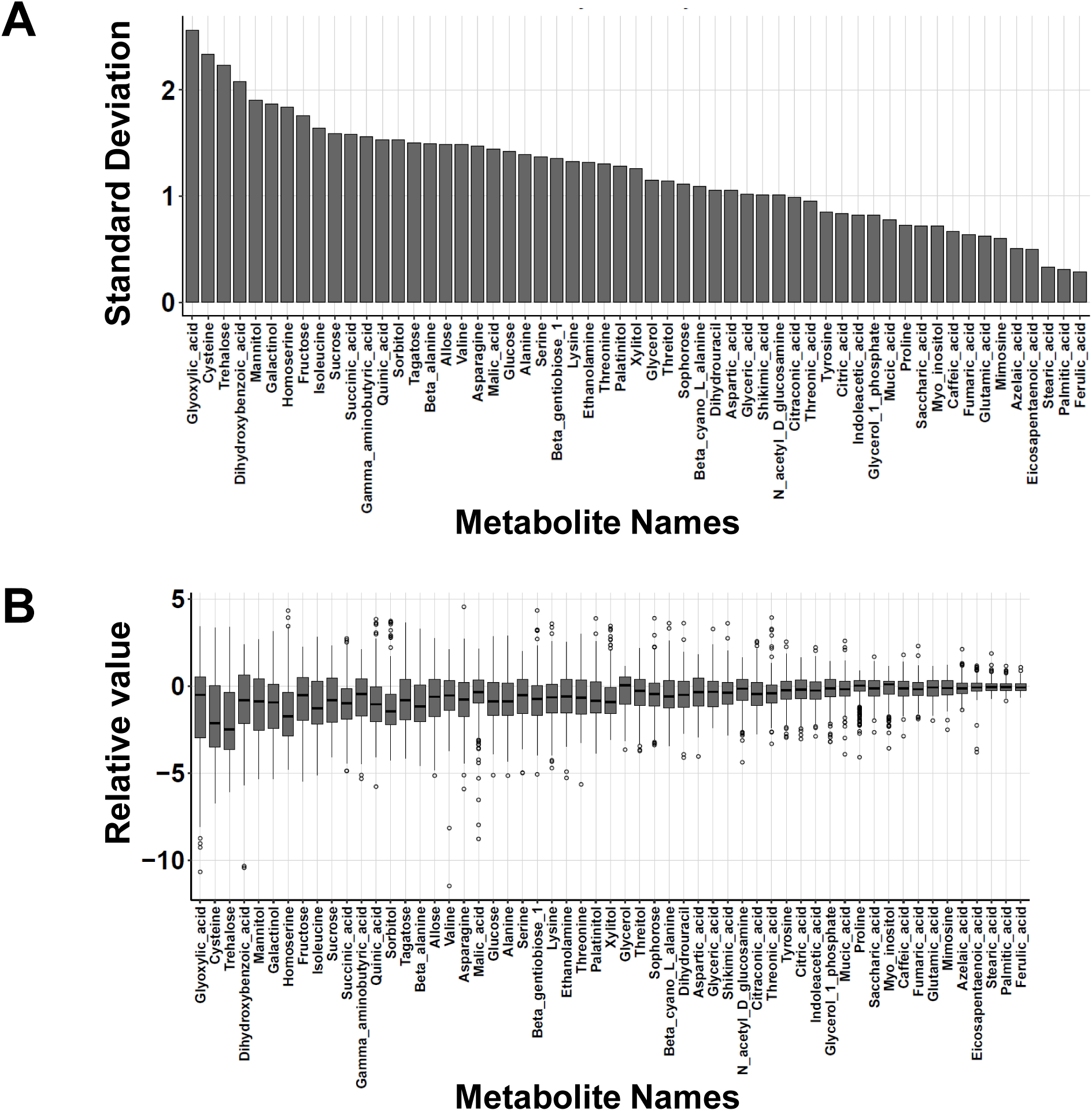
Metabolite level diversity in maize kernels. **(A)** Ranked standard deviation (SD) of 57 metabolite levels quantified across 265 maize inbred lines. Metabolites are sorted from highest variation (left) to lowest variation (right). **(B)** Box plots showing the distribution of relative abundance for each metabolite, arranged in the same order as in panel A. The central line represents the median, the box edges represent the interquartile range, and dots represent outliers.

### mGWAS identified 154 candidate genes associated with kernel metabolite levels

We performed mGWAS using a standard mixed linear model (Yu et al. 2006), and identified 6,039 significant locus–trait associations (*P* ≤ 6.5 ×10^-8^). To create a high-confidence list, we filtered these locus–trait associations based on signal density and statistical significance. First, we selected associations from genomic locations with signal counts exceeding 5% of the maximum number of signals (N) detected at a single location, yielding 212 associations. Second, we independently selected associations within the most significant 5% based on the *P* value, yielding 469 associations. The overlap of these two filtered sets resulted in 62 locus–trait associations. These associations spanned 20 metabolite traits and implicated 788 genes located within associated genomic intervals (Table S2). Among them, 154 (19.54%) were metabolic enzyme-coding genes (Table S3). Figure 2 shows representative Manhattan plots for associations involving enzyme-coding genes. Threonic acid and shikimic acid were significantly associated with SNPs located within a putative squalene synthase-encoding gene on chromosome 5 and near an anthocyanidin reductase-encoding gene on chromosome 10, respectively (Fig 2A and 2B). Similarly, sorbitol and citric acid were associated with loci on chromosomes 1 and 7, respectively, where the annotated candidate genes within the associated intervals encoded riboflavin synthase and peroxidase (Fig 2C and 2D). We also identified Zm00001d034245, which encodes a cytosolic purine 5-nucleotidase, as a potential pleiotropic enzyme-coding gene significantly associated with both 5,6-dihydroxyuracil and β-cyano-L-alanine levels (Fig 2E and 2F).

**Figure 2.**
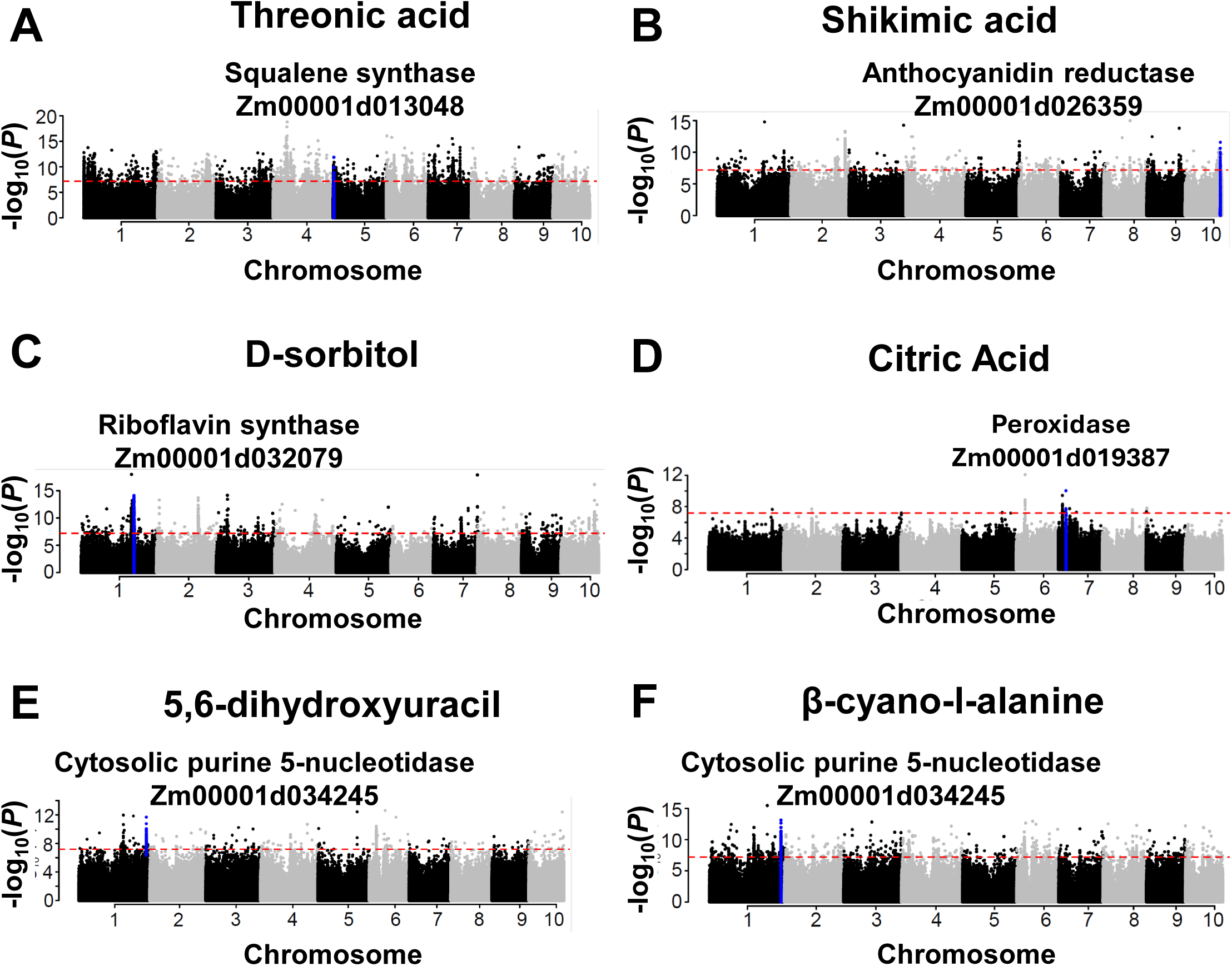
Genome-wide association mapping for kernel metabolite levels. The Manhattan plot displays the results of mGWAS for **(A)** Theronic acid, **(B)** Shikimic acid, **(C)** D-sorbitol, **(D)** Citric acid, **(E)** 5,6-dihydroxyuracil, and **(F)** β-cyano-L-alanine contents in maize kernels. The y-axis shows the-log_10_(p) value for each association, and the x-axis shows the genomic position across the 10 maize chromosomes. The dashed horizontal line indicates the Bonferroni-adjusted significance threshold (6.5 × 10^−8^). Significant SNPs are indicated in blue, and the enzyme gene names with corresponding metabolites.

### Natural variation in the *Shikimate dehydrogenase 1* (*Sad1*) gene is associated with homoserine levels in maize kernels

The mGWAS for homoserine accumulation revealed a major association peak on chromosome 10 (Figure 3A). The corresponding Q-Q plot showed a deviation from the expected distribution primarily at the tail, consistent with a robust association signal and adequate control of population structure (Figure 3B). To investigate the genetic basis of this signal, we analyzed the 2Mb locus, from 26Mb to 28Mb on chromosome 10, surrounding the most significant SNP (blue dot; Figure 3C). The regional Manhattan plot (Figure 3C) revealed a region of high linkage disequilibrium (LD), which contained approximately 600 significant SNPs and 32 gene models (Figure 3C, gray bars).

**Figure 3.**
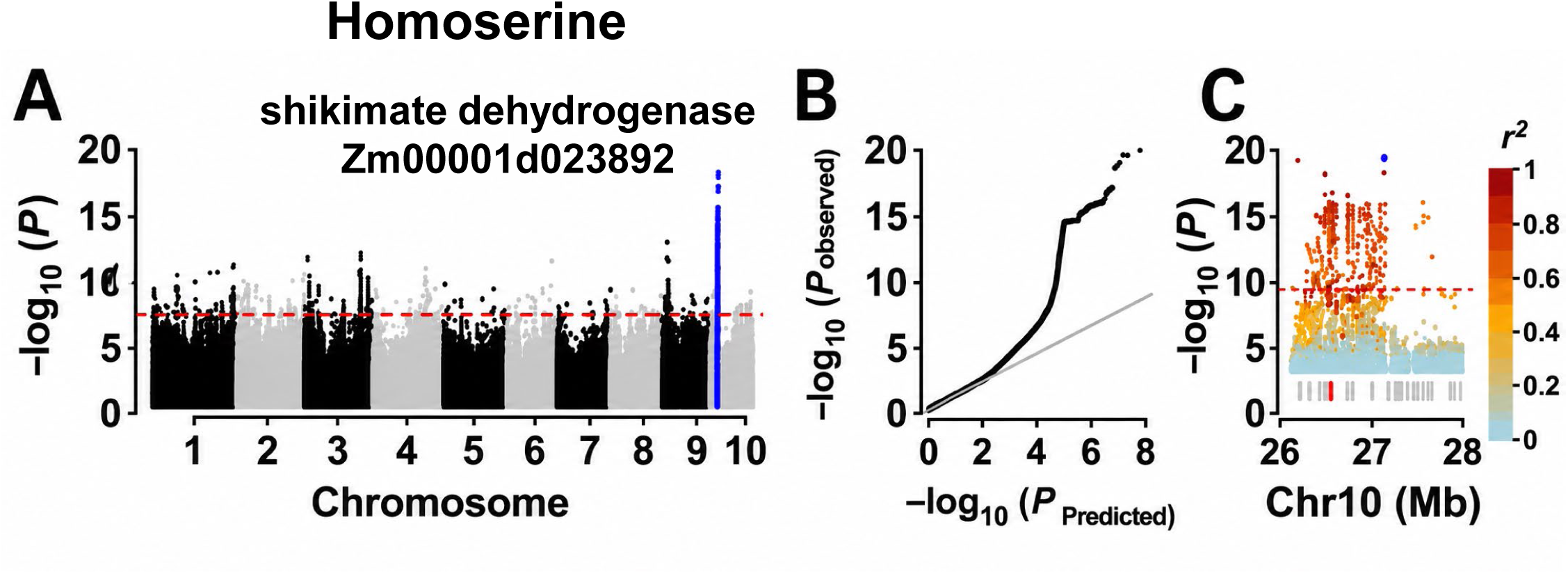
Genome-wide association mapping and regional analysis for kernel homoserine content. **(A)** A Manhattan plot for kernel homoserine contents. The y-axis shows the –log_10_(*P*) value for each association, and the x-axis shows the genomic position. The dots indicate individual SNPs. The dashed horizontal line indicates the Bonferroni-adjusted significance threshold (*P* = 6.5 × 10^−8^). SNPs at the primary association peak located on chromosome 10 are colored blue. **(B)** Quantile–quantile plot for the homoserine mGWAS result. The x-axis and y-axis represent −log_10_ transformed predicted p-values and observed p-values, respectively. The dots indicate individual SNPs and the diagonal line represents the expected values under the null hypothesis for no association. **(C)** A regional Manhattan plot for the 2Mb locus on chromosome 10. The dashed line represents the Bonferroni-adjusted significance threshold (*P* = 6.5 × 10^−8^). The lead SNP is shown with a blue dot. The other SNPs are colored according to their linkage disequilibrium (*r*^2^) with the lead SNP, as indicated by the scale bar with blue (0) to red (1). Gene models in the region are shown as light blue bars on the x-axis, with the location of the Shikimate Dehydrogenase 1 (sad1, Zm00001d023892) gene highlighted by a red bar.

Among these genes, we prioritized *Zm00001d023892*, which encodes Shikimate dehydrogenase 1 (*Sad1*), based on its predicted biological function and the presence of multiple significant polymorphisms in or near the gene. Shikimate dehydrogenase is a metabolic enzyme that potentially impacts the primary metabolic network involving homoserine (Wen et al. 2015). We identified 55 significant SNPs in or 10 kb upstream region of the *Sad1* gene. Of these, 37 were in the 1100 bp upstream region, and 18 were within the gene body, including four in coding regions. This combination of a strong statistical association at the locus, a plausible biological function, and multiple potentially functional polymorphisms within the gene made *Sad1* a promising candidate for functional analysis.

Association mapping and pairwise LD analysis on the *Sad1* coding region identified four significant SNPs within the *Sad1* coding region (Figure 4A). Two variants in exon 1 (SNP_26542548, *P* = 6.81 × 10⁻¹³; SNP_26542558, *P* = 4.03 × 10⁻¹²) and one in exon 9 (SNP_26546473, *P* = 6.33 × 10⁻¹³) and one in exon 10 (SNP_26546743, *P* = 3.94 × 10⁻¹²) were significantly associated with the kernel homoserine levels. LD analysis revealed that these exon variants were in high LD (*r²* > 0.9), suggesting that they are frequently inherited together as a functional unit. All 265 analyzed maize inbreds with reliable genome sequence data were classified into two haplotypes based on the combination of these four significant SNPs (Figure 4A). A large group of 213 inbred lines, including B73, carried Haplotype 1 (CCCA), whereas 19 inbred lines carried Haplotype 2 (GTTT); these lines belonged to the non-stiff stalk subpopulation (Table S4). Haplotype 2 inbreds showed significantly higher homoserine accumulation than Haplotype 1 (Figure 4B).

**Figure 4.**
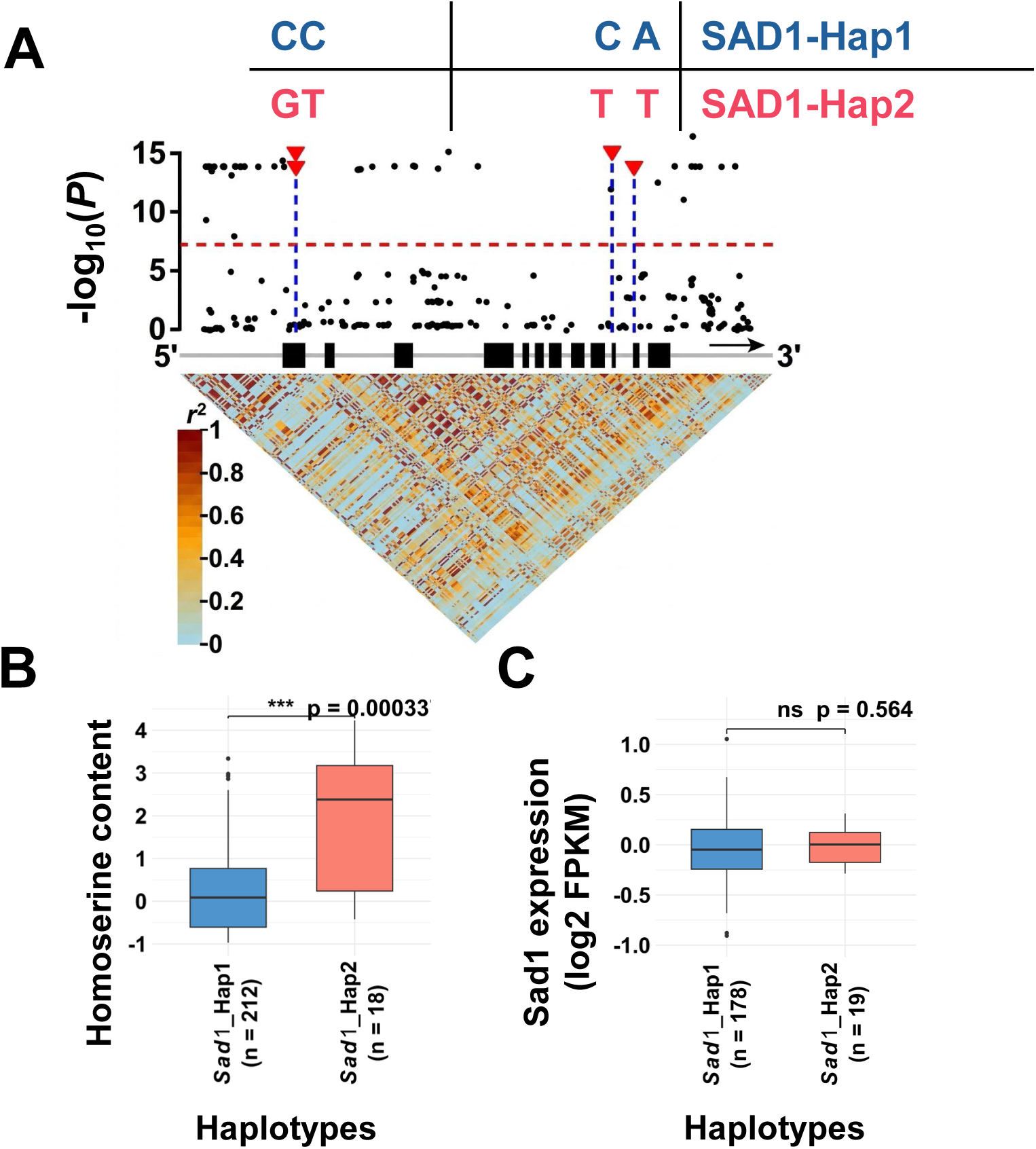
*Sad1* haplotypes and their association with homoserine content and gene expression. **(A)** Zm00001d023892 (*Sad1*) gene model and linkage disequilibrium (LD) plot. Black boxes represent exons. Black dots indicate the –log_10_(*P*) value for the association between each SNP and kernel homoserine levels in the mGWAS analysis shown in Figure 3. The four red triangles indicate significant SNPs in coding regions. The heatmap shows the pairwise LD (*r*^2^) values between SNPs in the region, with the inset table listing specific *r*^2^ values among the four SNPs in the coding regions. The two haplotypes formed by the four coding region SNPs. **(B)** Boxplot comparing homoserine contents between the two haplotype groups. The x-axis represents the haplotype and the number of inbreds belonging to each haplotype group in the analyzed population. The y-axis represents the log_2_-transformed relative homoserine levels. The difference was statistically significant (Student’s t-test, *p* < 0.001). **(C)** Boxplot comparing *Sad1* gene expression between the two haplotype groups. The x-axis represents the haplotype and the number of inbreds belonging to each haplotype group. The y-axis represents the log_2_ relative FPKM values of *Sad1* expression. There was no significant difference in gene expression levels (Student’s t-test, *p* = 0.564).

To determine whether these variants influence kernel homoserine levels via gene expression regulation, we analyzed homoserine accumulation alongside previously published *Sad1* gene expression data from developing kernels (Kremling et al. 2018). A regression analysis across the diversity panel revealed a very weak correlation between *Sad1* mRNA abundance and homoserine content (R^2^ = 0.10; Figure S1), indicating that expression variation explained only a minor fraction of the metabolic variance. Furthermore, *Sad1* expression did not differ significantly between the two haplotype groups (*P* =0.564; Table S5 and Figure 4C). Collectively, these data suggest that natural variation in the *Sad1* coding sequence likely influences shikimate dehydrogenase enzyme activity or stability rather than gene expression, affecting kernel homoserine accumulation.

### Non-synonymous SNPs alter SAD1 functional domains

To investigate the functional consequences of the identified *Sad1* haplotypes, we predicted the impact of coding-sequence polymorphisms on SAD1 protein structure. Of the four SNPs identified in the coding regions, three resulted in non-synonymous amino acid substitutions: leucine to phenylalanine at position 46 (L46F), proline to serine at position 522 (P522S), and glutamine to leucine at position 537 (Q537L; Figure 5A). Protein domain annotation mapped these substitutions to distinct functional domains of the enzyme. The L46F substitution is located within the N-terminal type I 3-dehydroquinase domain, while P522S and Q537L were located within the C-terminal NADP-binding domain (Figure 5A). The positions of these substitutions within catalytic and cofactor-binding domains suggest that they likely impact enzyme catalytic function, cofactor interaction, or local protein stability.

**Figure 5.**
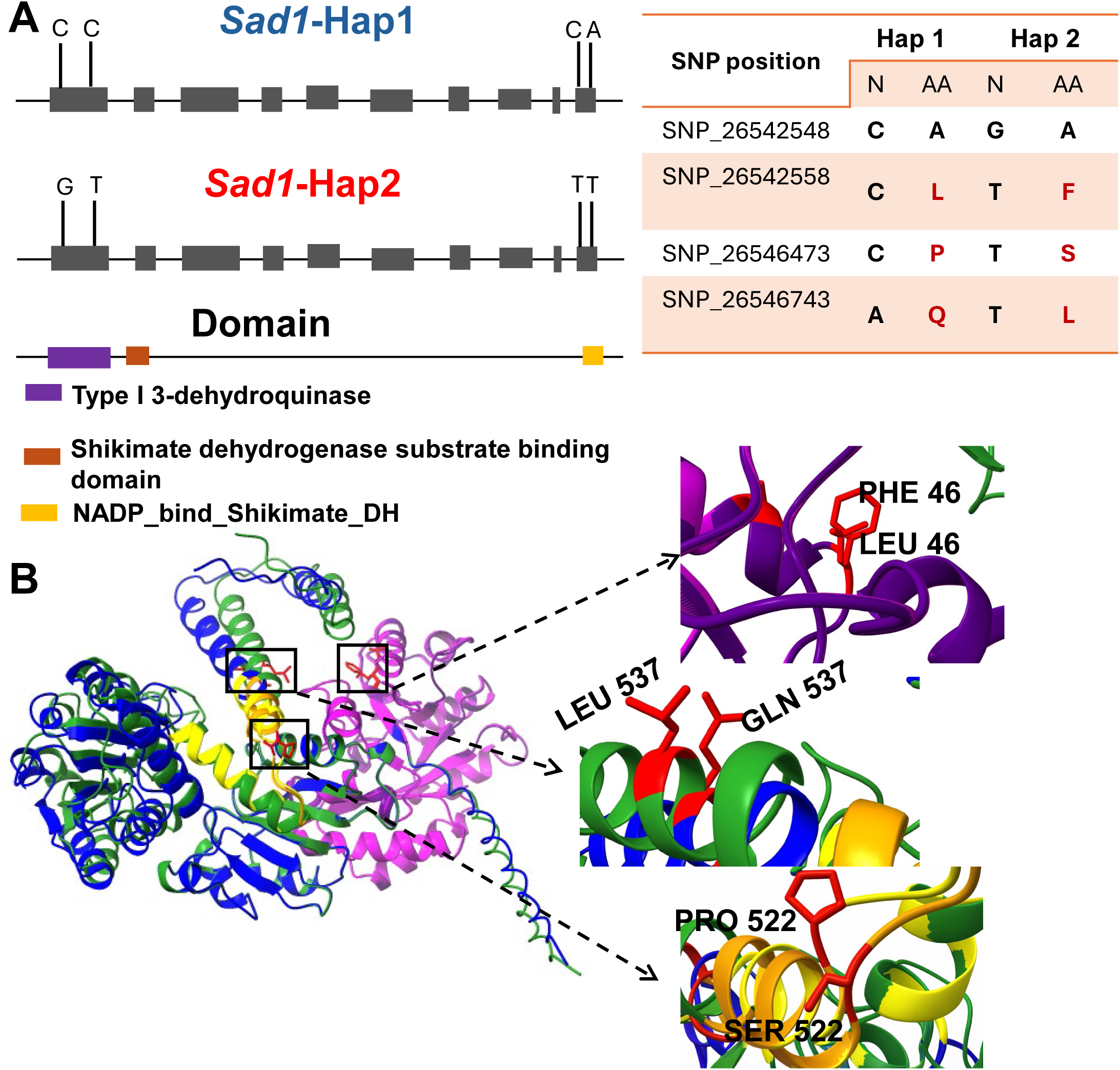
Predicted structural impacts of non-synonymous variants on the SAD1 enzyme protein. **(A)** Gene and protein domain architecture of SAD1 protein encoded by Zm00001d023892. The schematic maps the four coding-region SNPs associated with kernel homoserine levels to the exon structure (gray boxes) and the protein domains: the type I 3-dehydroquinase domain (purple), the shikimate dehydrogenase substrate binding domain (brown), and the NADP-binding shikimate dehydrogenase domain (orange). The table shows the SNP positions, their nucleotide (N), and the resulting amino acid (AA) between SAD1-Hap1 and SAD1-Hap2 sequences. Non-synonymous amino acid substitutions are highlighted in red. **(B)** AlphaFold protein structure predictions for the SAD1-Hap1 and SAD1-Hap2 are superimposed (left). The type I 3-dehydroquinase domain (purple/magenta) and the NADP-binding shikimate dehydrogenase domain (yellow/orange) are highlighted. Dashed lines indicate the regions enlarged on the right. The upper inset highlights the substitution of Leu46 to Phe46, the middle inset shows Gln537 to Leu 537 substitution, and the lower inset displays substitution of Pro522 to Ser522. Substituted residues are shown as sticks to illustrate their positions.

AlphaFold-based structural modeling of the two SAD1 haplotypes showed good backbone alignment, indicating a conserved global fold (Figure 5B). However, the specific side-chain substitutions suggest potential impacts on enzymatic properties. In the N-terminal domain, the L46F substitution replaces an aliphatic leucine with a bulky aromatic phenylalanine. The structural superposition suggested this volume increase might introduce steric hindrance or disrupt the hydrophobic packing essential for domain stability (Figure 5B). In the C-terminal domain, the substitution of rigid proline (P522S) and a polar glutamine (Q537L) within the NADP-binding pocket (Figure 5B) could alter cofactor interaction or local backbone flexibility. These structural predictions suggest that the haplotype variants likely modulate the overall enzymatic activity of SAD1.

### Genetic variation alters catalytic efficiency of shikimate dehydrogenase

To evaluate the effects of the structural variations in SAD1 enzyme on its catalytic function, we generated recombinant enzymes of SAD1 Haplotype 1 (SAD1-Hap1) and the variant containing L46F, P522S, and Q537L substitutions resembling Haplotype 2 (SAD1-Hap2; Figure S2). In the presence of saturating concentrations of NADP^+^ (1 mM) and shikimate (5 mM), SAD1-Hap1 exhibited substantially higher NADPH production (Figure 6A), resulting in a specific activity 13.48 times higher than SAD1-Hap2 (Figure 6B). Since two of the amino acid substitutions were located in the NADP^+^ binding domain (Figure 5), we determined the steady-state kinetic parameters of the haplotype enzymes by varying NADP+ concentrations (Figure 6C, Table 1). Enzyme kinetics characterization revealed that the *V*_max_ of SAD1-Hap1 was 23.7 ± 4.3 µmol min^-1^ which is about eleven-fold higher than that observed *V*_max_ for SAD1-Hap2 (2.1 ± 0.13 µmol min^-1^). This activity is reflected by the catalytic turnover rate (*k*_cat_) where SAD1-Hap1 achieved 0.168 ± 0.031 s^-1^ compared to 0.0149 ± 0.0009 s^-1^ for Hap2. Overall, SAD1-Hap2 has significantly lower catalytic efficiency but slightly higher NADP^+^ affinity than SAD1-Hap1.

**Figure 6.**
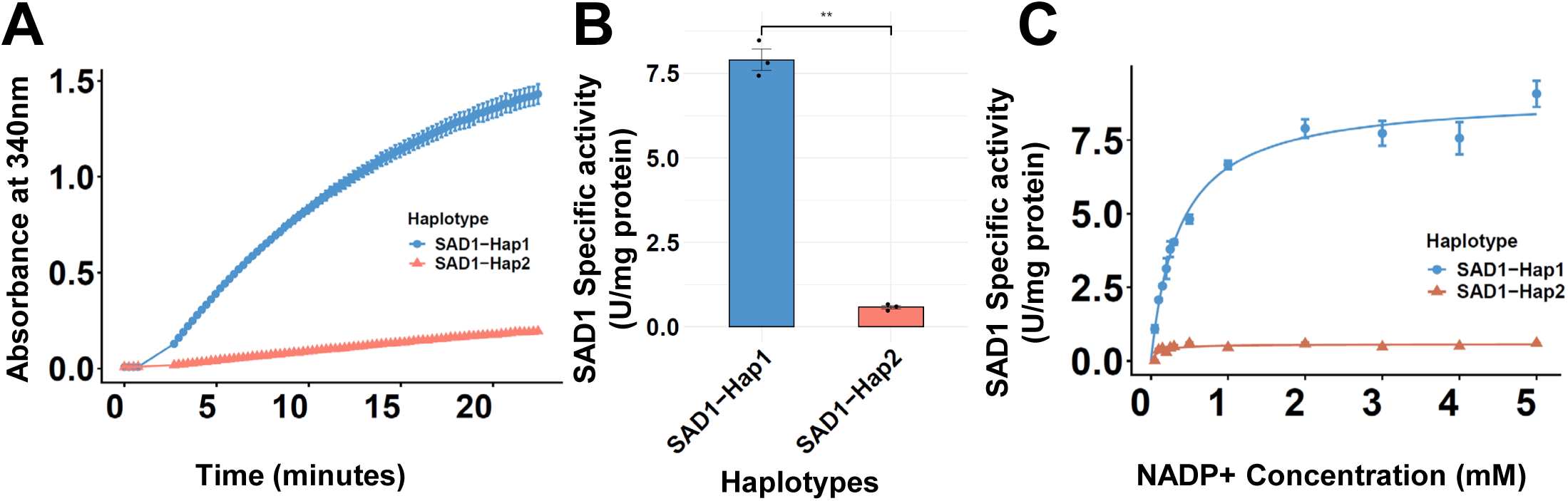
Enzyme kinetics of maize SAD1 haplotypes. **(A)** Time-course of shikimate dehydrogenase activity monitored by the increase in absorbance at 340 nm, reflecting NADPH formation. The activities of Haplotype 1 (SAD1-Hap1, blue) and Haplotype 2 (SAD1-Hap2, red) enzymes were measured. **(B)** Specific shikimate dehydrogenase activity at a NADP+ concentration of 2 mM. Bars represent mean specific activity (U mg⁻¹ protein) ± SE, with individual technical replicates shown as dots. Statistical significance was determined using an unpaired two-tailed Welch’s *t*-test (P < 0.0001). **(C)** NADP^+^-dependent enzyme activities of SAD1-Hap1 (blue) and SAD1-Hap2 (red) using NADP⁺ as the variable substrate. Data points represent mean specific activity (U mg⁻¹ protein) ± SE of three independent measurements, and curves represent nonlinear regression fits to the Michaelis–Menten equation.

**Table 1.** Kinetic parameters of SAD1-Hap1 and SAD1-Hap2 for NADP⁺.

| Protein | $V_{\max}$ ($\mu\text{molmin}^{-1}$) | $K_m$ ($\mu\text{M}$) | $k_{\text{cat}}$ ($\text{s}^{-1}$) |
| --- | --- | --- | --- |
| SAD1-Hap1 | $23.7 \pm 4.3$ | $241.0 \pm 30.0$ | $0.168 \pm 0.031$ |
| SAD1-Hap2 | $2.1 \pm 0.13$ | $131.4 \pm 3.0$ | $0.0149 \pm 0.0009$ |

### Metabolic network modeling suggested a connection between SAD1 activity and homoserine accumulation through redox-coupled oxaloacetate partitioning

To evaluate the influence of shikimate dehydrogenase activity on kernel homoserine accumulation, we used a genome-scale metabolic network model of maize kernels (Chowdhury et al., 2023). The reference model predicted the SAD1-mediated plastidial shikimate dehydrogenase flux of 0.105 mmol gDW^-1^ h^-1^. We progressively constrained this reaction to values as low as 0.011 mmol gDW^-1^ hour^-1^, one-tenth of the reference value, and estimated the resulting homoserine pool size. Decreasing SAD1 flux produced a linear increase in predicted homoserine pool size (Figure 7A). This relationship was consistent with the greater homoserine accumulation observed in inbreds carrying SAD1-Hap2, whose encoded enzyme exhibited lower catalytic efficiency (Figures 4 and 6).

**Figure 7.**
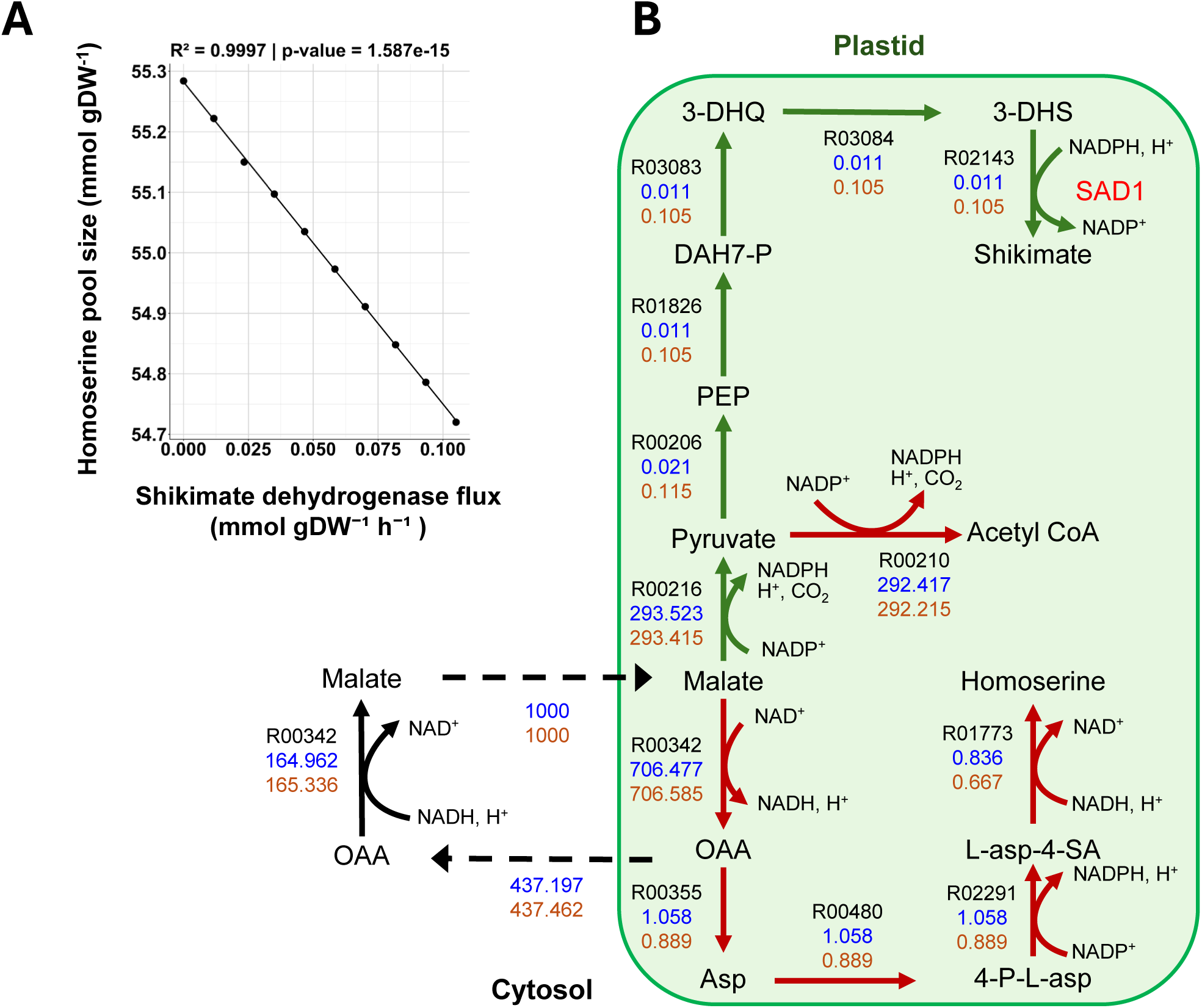
Flux balance simulation of homoserine accumulation and metabolic flux redistribution in response to altered plastidial SAD1 reaction flux.(A) Predicted homoserine pool size when the plastidial SAD1-catalyzed shikimate dehydrogenase reaction was progressively constrained from the reference flux of 0.105 mmol gDW⁻¹ h⁻¹ to 0.011 mmol gDW⁻¹ h⁻¹. Each point represents predicted homoserine pool size (mmol gDW^-1^). The line shows a linear regression fit (R² = 0.9997, *P* = 1.587 × 10⁻¹⁵). (B) Flux distribution through reactions connecting plastidial shikimate metabolism and the aspartate/homoserine biosynthetic pathway. Values adjacent to each reaction indicate KEGG reaction identifiers and the predicted fluxes under the lowest SAD1 flux condition of 0.011 mmol gDW⁻¹ h⁻¹ (blue) and the reference SAD1 flux condition of 0.105 mmol gDW⁻¹ h⁻¹ (brown). Green arrows indicate reaction fluxes that were positively correlated with SAD1 flux, whereas dark red arrows indicate reaction fluxes that are negatively correlated with SAD1 flux. Blue numbers represent reaction fluxes under the lower shikimate dehydrogenase flux condition, and brown numbers represent reaction fluxes under the higher shikimate dehydrogenase flux condition. Dashed arrows indicate metabolite transport between the cytosol and plastid. OAA, oxaloacetate; PEP, phosphoenolpyruvate; Asp, aspartate; 4-P-L-asp, 4-phospho-L-aspartate; L-asp-4-SA, L-aspartate-4-semialdehyde; DAH7-P, 3-deoxy-D-arabino-heptulosonate-7-phosphate; 2-DHQ, 2-dehydroquinate; 3-DHS, 3-dehydroshikimate.

To reveal metabolic pathway alterations underlying this relationship, we identified plastidial reactions whose fluxes covaried with SAD1 flux (Table S6) and mapped them onto the relevant pathways (Figure 7B). Constraining SAD1 flux to lower values reduced shikimate pathway reactions from pyruvate to shikimate (R00206, R01826, R03083, R03084, and R02143) to similar extents, indicating that SAD1 flux restricted the overall pathway flux. Lower SAD1 flux was also associated with decreased plastidial malate dehydrogenase flux (R00342) and reduced oxaloacetate export to the cytosol (C00036). In parallel, flux through plastidial NADP^+^-dependent malic enzyme (R00216) and pyruvate dehydrogenase (R00210) increased slightly, indicating modestly greater allocation of malate-derived carbon to acetyl-CoA production. Reduced OAA export was accompanied by increased flux through the aspartate-derived homoserine pathway, including reactions catalyzed by aspartate aminotransferase (R00355), aspartate kinase (R00480), aspartate-semialdehyde dehydrogenase (R02291), and homoserine dehydrogenase (R01773). Together, the model indicated a mechanism in which reduced SAD1 flux alters plastidial carbon partitioning, decreases oxaloacetate export, and favors the allocation of plastidial oxaloacetate to homoserine biosynthesis.

## Discussion

The metabolite composition of maize grain is a crucial determinant of its nutritional and economic value. While previous studies have highlighted the extensive biochemical diversity of kernels of maize germplasm (Zhou et al. 2019) with diverse genetic architectures (Mural et al. 2022), understanding the genetic basis of this variation remains a key challenge for crop improvement. Our metabolite profiling of the diversity panel revealed a spectrum of variability in metabolite contents in kernels (Figure 1, Supplementary Figure 1). We observed that a subset of amino acids and sugar derivatives, most notably homoserine and trehalose, exhibited massive variation. Interestingly, the high metabolic variability did not always translate to significant genomic associations. This aligns with previous studies suggesting that primary metabolic traits are often governed by a highly polygenic architecture and are subject to significant environmental plasticity, making individual genetic drivers difficult to isolate (Keurentjes and Sulpice 2009; Wen et al. 2015). Whereas metabolites such as homoserine appear to be controlled by large-effect loci (Figures 2 and 3), other variable metabolites are likely governed by polygenic architecture with many small-effect genes or possess lower heritability due to environmental plasticity. Thus, while metabolic diversity is a prerequisite for genetic mapping, the discovery of specific causal genes relies on the presence of major functional alleles segregating within the population. Thus, the discovery of *Sad1* represents a distinct case where a specific, large-effect coding variant drives a major shift in primary metabolism.

We revealed that the genetic variation in a shikimate dehydrogenase gene (*Sad1*) is associated with homoserine accumulation in maize kernels, suggesting the influence of SAD1 activity on aspartate family amino acid metabolism (Figure 3). Our results suggest that the naturally occurring haplotypes within the *Sad1* gene coding regions are key contributors to the variation in homoserine accumulation observed in maize. The kernel *Sad1* gene expression and homoserine contents did not show correlation (Figure 4C, Figure S1), suggesting that the functional diversity of the SAD1 enzyme affects homoserine accumulation. The four SNPs within the coding regions showed a strong linkage disequilibrium, leading to two haplotypes distinct in kernel homoserine levels in the tested inbreds (Figure 4). A co-inheritance of SNPs in coding sequences has been reported for an *O-methyltransferase* gene in rice, which leads to haplotypes with distinct trigonelline levels in leaves (Chen et al. 2014). Unlike the SNPs in the rice methyltransferase, the co-inherited SNPs in *Sad1* span exon regions encoding distant functional domains (Figure 5A). The tight linkage between these distant residues suggests they may be functionally co-adapted. Natural selection likely preserved this specific combination because the mutations work synergistically to maintain enzyme stability or kinetic balance, whereas intermediate combinations may have been functionally compromised.

Our structural modeling showed that the non-synonymous SNPs distinguishing the haplotypes cause amino acid alterations within the catalytic and cofactor-binding domains of the SAD1 enzyme (Figure 5). These amino acid alterations enhanced NADP^+^ affinity but impaired the catalytic efficiency of SAD1-Hap2 enzyme (Table 1, Figure 6). This uncoupling of binding and catalysis may be indicative of a “thermodynamic pit,” where mutations inadvertently over-stabilize a non-productive ground-state complex (Bar-Even et al. 2011; Sanchez-Ruiz 2025), which is reported in various enzymes (Hegazy and Richard 2023). Alternatively, the co-inheritance of residue substitutions in dehydroquinate dehydrogenase and shikimate dehydrogenase domains may affect substrate channeling, which is reported in the homologous enzyme in *Arabidopsis thaliana* (Singh and Christendat 2006). However, further studies are required to elucidate the precise mechanisms underlying the differences in kinetic characteristics between SAD1-Hap1 and SAD1-Hap2.

The simulated negative correlation between SAD1-mediated plastidial shikimate dehydrogenase flux and homoserine pool size (Figure 7A) supports a quantitative connection between SAD1 enzyme activity and kernel homoserine content. This relationship is consistent with the biochemical properties of the SAD1 haplotypes. SAD1-Hap2 enzyme exhibited substantially reduced catalytic efficiency compared with SAD1-Hap1 (Figure 6, Table 1), and inbreds harboring Haplotype 2 enzymes accumulated higher levels of homoserine (Figure 4B). These results suggest that reduced SAD1 activity in Haplotype 2 inbreds likely alters metabolic flux favoring homoserine accumulation.

Our pathway analysis further suggests that the negative relationship between SAD1 activity and homoserine biosynthesis is likely mediated by changes in plastid redox metabolism and carbon partitioning through the malate-oxaloacetate shuttle. SAD1 catalyzes an NADPH-consuming reaction in the plastid, creating a demand for reducing equivalents. Since the non-photosynthetic kernel plastids lack the ability to produce NADPH via photosynthesis, the NADPH supply depends on heterotrophic metabolism (Selinski and Scheibe 2019). In the metabolic network simulation, fluxes through eight reactions involving redox cofactors covaried with SAD1 flux (Table S6), suggesting that altered SAD1 activity is coupled to broader changes in plastidial redox-associated metabolism.

Notably, the model predicted that flux through part of the malate–oxaloacetate shuttle (R00342 and C00036) was positively associated with SAD1 flux (Figure 7B). In this context, the shuttle can transfer reducing equivalents from the cytosol to the plastid (Selinski and Scheibe 2019) to support the NADPH-dependent SAD1 reaction. At the same time, this exchange can affect the partitioning of oxaloacetate between the shuttle and amino acid biosynthesis. Increased SAD1 flux was associated with enhanced malate–oxaloacetate shuttle activity and oxaloacetate export from the plastid, thereby reducing the availability of plastidial oxaloacetate for aspartate biosynthesis and downstream homoserine production (Figure 7B). Conversely, reduced SAD1 flux in SAD1-Hap2-harboring inbreds is predicted to lower the demand for imported reducing equivalents, decrease flux through this shuttle, and retain more oxaloacetate for aspartate-family amino acid biosynthesis, resulting in increased homoserine accumulation. Collectively, the metabolic network modeling suggests that variation in SAD1-mediated shikimate pathway flux potentially influences plastidial carbon metabolism through coordinated reorganization of redox metabolism and carbon partitioning.

In summary, this study demonstrates that natural variation in the *Sad1* coding sequence is a key determinant of homoserine levels in maize kernels. Three linked non-synonymous SNPs are inherited together as a unit, forming two distinct haplotypes that produce enzyme variants with distinct catalytic efficiency that likely influence plastidial redox metabolism and oxaloacetate partitioning to the aspartate/homoserine biosynthesis pathways. This metabolic balancing probably affects the accumulation of homoserine in maize kernels. Notably, the Haplotype 2 variant, which is associated with higher homoserine levels, is present at low frequency in the diversity panel, suggesting that maintaining high flux through the shikimate pathway and following phenylpropanoid pathways may be favored through the breeding process, likely due to biosynthesis of lignin and defense compounds (Tohge et al., 2013).

These findings identify the malate–oxaloacetate shuttle as the mechanistic link connecting the shikimate and homoserine pathways and demonstrate that changes in secondary metabolism can indirectly influence primary amino acid biosynthesis through plastid redox balance. This work provides new fundamental insight into the metabolic coordination between specialized and primary metabolism and highlights the importance of redox-mediated metabolic integration in plants. Additionally, this study showed that an innovative integration of population genetics, structural enzymology, and systems-level metabolic modeling establishes a causal framework connecting natural enzyme variation to metabolite network regulation. The application of metabolic flux simulation using a metabolic network model to the implication of GWAS-based association analysis is particularly useful to evaluate the mechanistic plausibility of genetic associations.

## Author contributions

Experimental concept and design: T.O., J.Y., and R.H.; Field experiment design and sample collection: J.Y.; Metabolite extraction and QTOF GC-MS run: C.P.; Perform Genome wide association study: G.X.; Data analysis and interpretation: R.H.; Vector construction: R.H.; Cloning, gene expression and protein purification: T.D.; Enzyme kinetics assay: R.H. and T.D.; Metabolite network model development and simulation: N.B.C. and R.S.; Manuscript writing and figures: R.H. and T.O.; Manuscript review: all authors read and approved the final manuscript.

## Funding

This work was supported by the United States Department of Agriculture National Institute of Food and Agriculture (USDA-NIFA: 2021-67013-33898 and NEB-30-134) to T.O and J.Y.

## Conflicts of interest

None declared.

## Supporting information

Supplementary doc 1

## Data availability

*Sad1* gene expression data from (Kremling et al. 2018) were used in this study. All codes used to generate maize kernel metabolite network model were in the following Github repository: https://github.com/niazchebuet/maize_GWAS.

## Supplementary Materials

**Supplementary Table 1.** Metabolite abundance data for 57 metabolites across 265 maize inbred lines.

**Supplementary Table 2.** Candidate genes associated with significant metabolite genome-wide association study (mGWAS) loci.

**Supplementary Table 3.** Metabolic enzyme-coding genes among mGWAS candidate genes.

**Supplementary Table 4.** *Sad1* haplotype assignments and kernel homoserine contents of maize inbred lines.

**Supplementary Table 5.** SAD1 haplotype assignments and SAD1 expression levels in maize inbred lines.

**Supplementary Table 6.** Predicted metabolic flux distributions under different constraints on plastidial SAD1 reaction flux.

**Supplementary doc 1.** pET21b with *Sad1*_Hap1 and pET21b with *Sad1*_Hap2 sequence

---

**Supplementary Figure 1.**
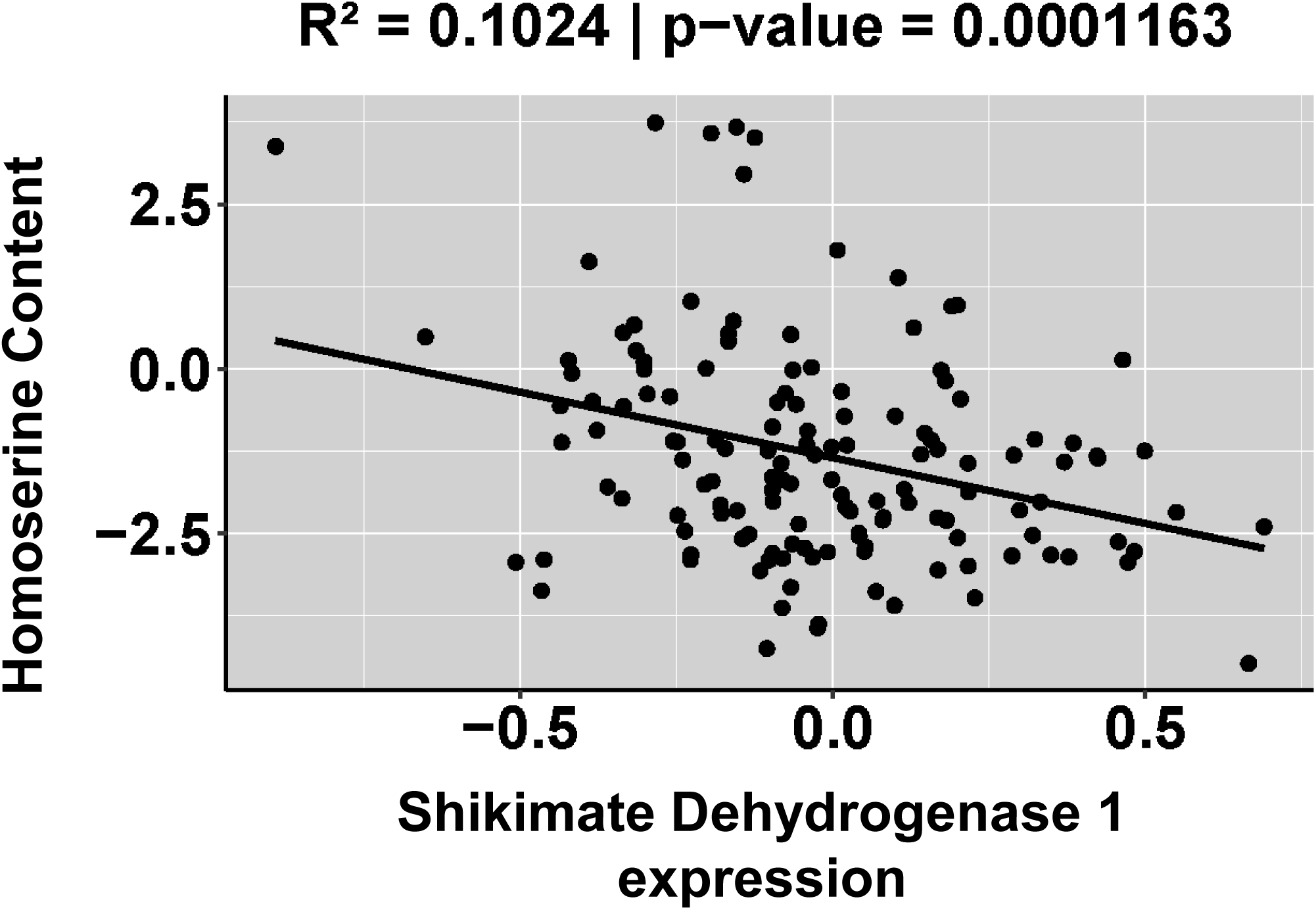
Correlation analysis between *Sad1* gene expression and kernel homoserine content. Scatter plot displaying the relationship between normalized *Sad1* transcript abundance (FPKM) and homoserine content across the diversity panel. The line represents the linear regression fit.

**Supplementary Figure 2.**
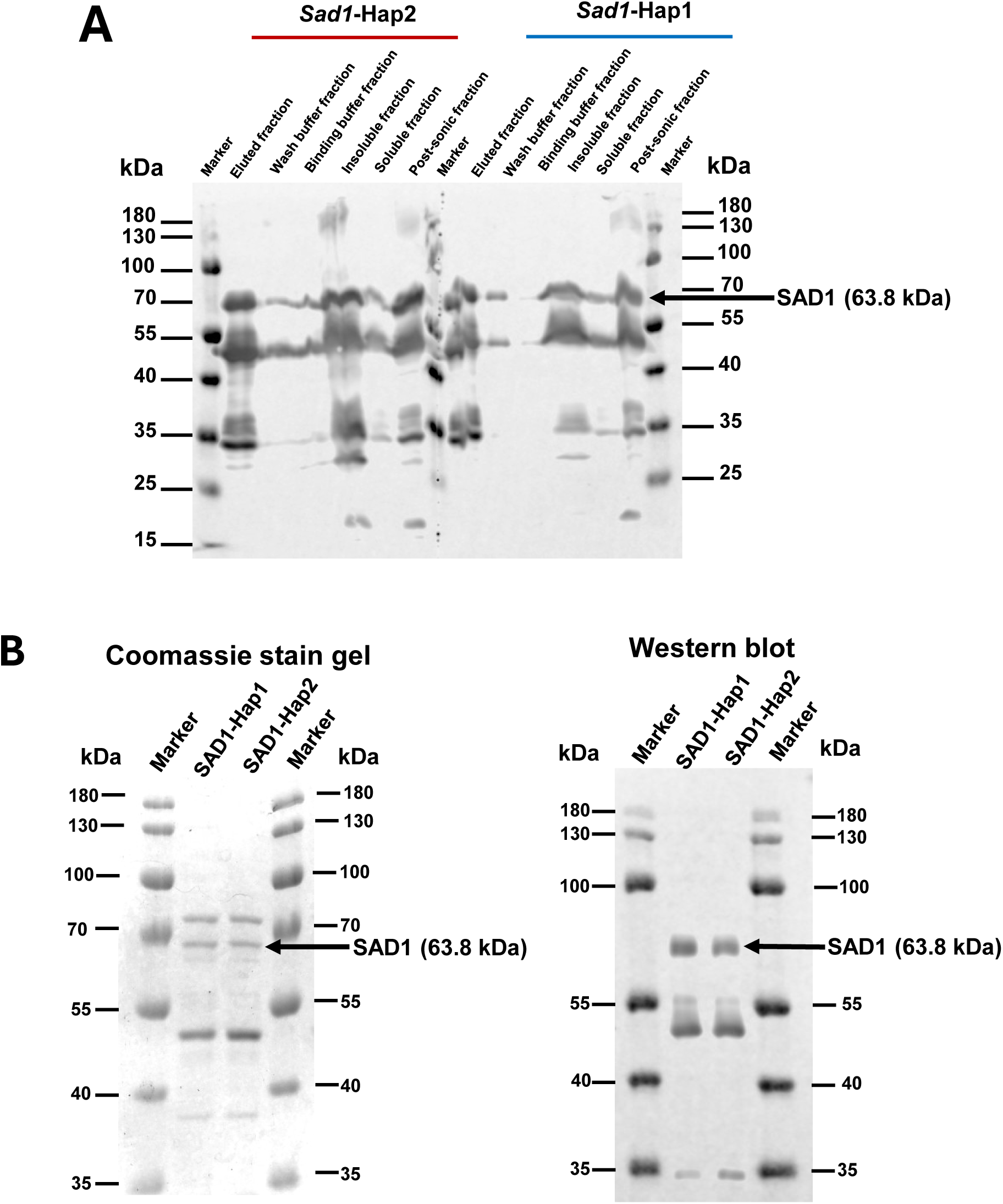
Expression and purification of recombinantSAD1-Hap1 and SAD1-Hap2 proteins using *Escherichia coli* BL21 cells. **(A)** Western-blotting analysis of different fractions collected during purification of recombinant SAD1-Hap1 and SAD1-Hap2 proteins, including elution, wash, binding, insoluble, soluble, post-lysis, and whole-cell (post-sonic) fractions. SAD1 proteins were detected using anti-HA antibody. Protein bands migrated to the expected SAD1 molecular weight of 63.8 kDa are indicated by an arrow. **(B)** Coomassie Brilliant Blue–stained SDS–PAGE gel (left) and anti-HA Western blot (right) of partially purified SAD1-Hap1 and SAD1-Hap2 proteins loaded at a total protein concentration of 7.5 µg mL^-1^.

## Notes

### Competing Interest Statement

The authors have declared no competing interest.

## References

Almeida AM, Abrahão J, Seixas FAV, Bueno PSA, Oliveira MASD, Tomazini LF, Constantin RP, Dos Santos WD, Marchiosi R, and Ferrarese-Filho O. Unraveling Shikimate Dehydrogenase Inhibition by 6-Nitroquinazoline-2,4-diol and Its Impact on Soybean and Maize Growth. Agronomy. 2024:14(5):930. 10.3390/agronomy14050930

Bar-Even A, Noor E, Savir Y, Liebermeister W, Davidi D, Tawfik DS, and Milo R. The Moderately Efficient Enzyme: Evolutionary and Physicochemical Trends Shaping Enzyme Parameters. Biochemistry. 2011:50(21):4402–4410. 10.1021/bi2002289

Bradbury PJ, Zhang Z, Kroon DE, Casstevens TM, Ramdoss Y, and Buckler ES. TASSEL: software for association mapping of complex traits in diverse samples. Bioinformatics. 2007:23(19):2633–2635. 10.1093/bioinformatics/btm308

Chang CC, Chow CC, Tellier LC, Vattikuti S, Purcell SM, and Lee JJ. Second-generation PLINK: rising to the challenge of larger and richer datasets. Gigascience. 2015:4(1):s13742–015-0047–8. 10.1186/s13742-015-0047-8

Chen J, Hu X, Shi T, Yin H, Sun D, Hao Y, Xia X, Luo J, Fernie AR, He Z, et al.Metabolite-based genome-wide association study enables dissection of the flavonoid decoration pathway of wheat kernels. Plant Biotechnol J. 2020:18(8):1722–1735. 10.1111/pbi.13335

Chen W, Gao Y, Xie W, Gong L, Lu K, Wang W, Li Y, Liu X, Zhang H, Dong H, et al. Genome-wide association analyses provide genetic and biochemical insights into natural variation in rice metabolism. Nat Genet. 2014:46(7):714–721. 10.1038/ng.3007

Chowdhury NB, Schroeder WL, Sarkar D, Amiour N, Quilleré I, Hirel B, Maranas CD, and Saha R. Dissecting the metabolic reprogramming of maize root under nitrogen-deficient stress conditions. J Exp Bot. 2022:73(1):275–291. 10.1093/jxb/erab435

Chowdhury NB, Simons-Senftle M, Decouard B, Quillere I, Rigault M, Sajeevan KA, Acharya B, Chowdhury R, Hirel B, Dellagi A, et al. A multi-organ maize metabolic model connects temperature stress with energy production and reducing power generation. iScience. 2023:26(12):108400. 10.1016/j.isci.2023.108400

Clark TJ, Guo L, Morgan J, and Schwender J. Modeling Plant Metabolism: From Network Reconstruction to Mechanistic Models. Annu Rev Plant Biol. 2020:71(1):303–326. 10.1146/annurev-arplant-050718-100221

Díaz-Quiroz DC, Cardona-Félix CS, Viveros-Ceballos JL, Reyes-González MA, Bolívar F, Ordoñez M, and Escalante A. Synthesis, biological activity and molecular modelling studies of shikimic acid derivatives as inhibitors of the shikimate dehydrogenase enzyme of *Escherichia coli*. J Enzyme Inhib Med Chem. 2018:33(1):397–404. 10.1080/14756366.2017.1422125

Du X, Huang G, He S, Yang Z, Sun G, Ma X, Li N, Zhang X, Sun J, Liu M, et al. Resequencing of 243 diploid cotton accessions based on an updated A genome identifies the genetic basis of key agronomic traits. Nat Genet. 2018:50(6):796– 802. 10.1038/s41588-018-0116-x

Flint-Garcia SA, Thuillet A, Yu J, Pressoir G, Romero SM, Mitchell SE, Doebley J, Kresovich S, Goodman MM, and Buckler ES. Maize association population: a high-resolution platform for quantitative trait locus dissection. Plant J. 2005:44(6):1054–1064. 10.1111/j.1365-313X.2005.02591.x

Giang VQ, Ky H, Thanh Tung NC, Hien NL, Van Manh N, Thanh NN, Thanh VC, and Yeap SK. Novel Deletion in Exon 7 of Betaine Aldehyde Dehydrogenase 2 (BADH2). Rice Sci. 2023:30(2):104–112. 10.1016/j.rsci.2023.01.003

Gomes De Oliveira Dal’Molin C, Quek L-E, Saa PA, Palfreyman R, and Nielsen LK. From reconstruction to C4 metabolic engineering: A case study for overproduction of polyhydroxybutyrate in bioenergy grasses. Plant Sci. 2018:273:50–60. 10.1016/j.plantsci.2018.03.027

Grafahrend-Belau E, Schreiber F, Koschützki D, and Junker BH. Flux Balance Analysis of Barley Seeds: A Computational Approach to Study Systemic Properties of Central Metabolism. Plant Physiol. 2009:149(1):585–598. 10.1104/pp.108.129635

Harrigan GG, Stork LG, Riordan SG, Reynolds TL, Ridley WP, Masucci JD, MacIsaac S, Halls SC, Orth R, Smith RG, et al. Impact of Genetics and Environment on Nutritional and Metabolite Components of Maize Grain. J Agric Food Chem. 2007:55(15):6177–6185. 10.1021/jf070494k

Hegazy R and Richard JP. Triosephosphate Isomerase: The Crippling Effect of the P168A/I172A Substitution at the Heart of an Enzyme Active Site. Biochemistry. 2023:62(20):2916–2927. 10.1021/acs.biochem.3c00414

Herrmann KM. The Shikimate Pathway: Early Steps in the Biosynthesis of Aromatic Compounds. Plant Cell. 1995:907–919. 10.1105/tpc.7.7.907

Johnson SR, Lange I, Srividya N, and Lange BM. Bioenergetics of Monoterpenoid Essential Oil Biosynthesis in Nonphotosynthetic Glandular Trichomes. Plant Physiol. 2017:175(2):681–695. 10.1104/pp.17.00551

Keurentjes JJB and Sulpice R. The role of natural variation in dissecting genetic regulation of primary metabolism. Plant Signal Behav. 2009:4(3):244–246. 10.4161/psb.4.3.7956

Kind T, Wohlgemuth G, Lee DY, Lu Y, Palazoglu M, Shahbaz S, and Fiehn O. FiehnLib: Mass Spectral and Retention Index Libraries for Metabolomics Based on Quadrupole and Time-of-Flight Gas Chromatography/Mass Spectrometry. Anal Chem. 2009:81(24):10038–10048. 10.1021/ac9019522

Kremling KAG, Chen S-Y, Su M-H, Lepak NK, Romay MC, Swarts KL, Lu F, Lorant A, Bradbury PJ, and Buckler ES. Dysregulation of expression correlates with rare-allele burden and fitness loss in maize. Nature. 2018:555(7697):520–523. 10.1038/nature25966

Lakshmanan M, Lim S-H, Mohanty B, Kim JK, Ha S-H, and Lee D-Y. Unraveling the light-specific metabolic and regulatory signatures of rice through combined in silico modeling and multi-omics analysis. Plant Physiol. 2015:pp.01379.2015. 10.1104/pp.15.01379

Lakshmanan M, Zhang Z, Mohanty B, Kwon J-Y, Choi H-Y, Nam H-J, Kim D-I, and Lee D-Y. Elucidating Rice Cell Metabolism under Flooding and Drought Stresses Using Flux-Based Modeling and Analysis. Plant Physiol. 2013:162(4):2140– 2150. 10.1104/pp.113.220178

Leemhuis H, Kelly RM, and Dijkhuizen L. Engineering of cyclodextrin glucanotransferases and the impact for biotechnological applications. Appl Microbiol Biotechnol. 2010:85(4):823–835. 10.1007/s00253-009-2221-3

Li Q, Yang X, Xu S, Cai Y, Zhang D, Han Y, Li L, Zhang Z, Gao S, Li J, et al. Genome-Wide Association Studies Identified Three Independent Polymorphisms Associated with α-Tocopherol Content in Maize Kernels. PLoS ONE. 2012:7(5):e36807. 10.1371/journal.pone.0036807

Maeda H and Dudareva N. The Shikimate Pathway and Aromatic Amino Acid Biosynthesis in Plants. Annu Rev Plant Biol. 2012:63(1):73–105. 10.1146/annurev-arplant-042811-105439

Mural RV, Sun G, Grzybowski M, Tross MC, Jin H, Smith C, Newton L, Andorf CM, Woodhouse MR, Thompson AM, et al. Association mapping across a multitude of traits collected in diverse environments in maize. GigaScience. 2022:11:giac080. 10.1093/gigascience/giac080

Orth JD, Thiele I, and Palsson BØ. What is flux balance analysis? Nat Biotechnol. 2010:28(3):245–248. 10.1038/nbt.1614

Palali Delen S, Xu G, Velazquez-Perfecto J, and Yang J. Estimating the genetic parameters of yield-related traits under different nitrogen conditions in maize. Genetics. 2023:223(4):iyad012. 10.1093/genetics/iyad012

Poolman MG, Kundu S, Shaw R, and Fell DA. Responses to Light Intensity in a Genome-Scale Model of Rice Metabolism. Plant Physiol. 2013:162(2):1060– 1072. 10.1104/pp.113.216762

Pott DM, Osorio S, and Vallarino JG. From Central to Specialized Metabolism: An Overview of Some Secondary Compounds Derived From the Primary Metabolism for Their Role in Conferring Nutritional and Organoleptic Characteristics to Fruit. Front Plant Sci. 2019:10:835. 10.3389/fpls.2019.00835

Riedelsheimer C, Lisec J, Czedik-Eysenberg A, Sulpice R, Flis A, Grieder C, Altmann T, Stitt M, Willmitzer L, and Melchinger AE. Genome-wide association mapping of leaf metabolic profiles for dissecting complex traits in maize. Proc Natl Acad Sci. 2012:109(23):8872–8877. 10.1073/pnas.1120813109

Saha R, Suthers PF, and Maranas CD. Zea mays iRS1563: A Comprehensive Genome-Scale Metabolic Reconstruction of Maize Metabolism. PLoS ONE. 2011:6(7):e21784. 10.1371/journal.pone.0021784

Sanchez-Ruiz JM. Binding *versus* catalysis in *de novo* enzyme design. Biocatal Biotransformation. 2025:1–4. 10.1080/10242422.2025.2577182

Selinski J and Scheibe R. Malate valves: old shuttles with new perspectives. Plant Biol. 2019:21(S1):21–30. 10.1111/plb.12869

Shende VV, Bauman KD, and Moore BS. The shikimate pathway: gateway to metabolic diversity. Nat Prod Rep. 2024:41(4):604–648. 10.1039/D3NP00037K

Singh SA and Christendat D. Structure of *Arabidopsis* Dehydroquinate Dehydratase-Shikimate Dehydrogenase and Implications for Metabolic Channeling in the Shikimate Pathway, Biochemistry. 2006:45(25):7787–7796. https://doi.org/10.1021/bi060366+

Skogerson K, Harrigan GG, Reynolds TL, Halls SC, Ruebelt M, Iandolino A, Pandravada A, Glenn KC, and Fiehn O. Impact of Genetics and Environment on the Metabolite Composition of Maize Grain. J Agric Food Chem. 2010:58(6):3600–3610. 10.1021/jf903705y

Wang W, Mauleon R, Hu Z, Chebotarov D, Tai S, Wu Z, Li M, Zheng T, Fuentes RR, Zhang F, et al. Genomic variation in 3,010 diverse accessions of Asian cultivated rice. Nature. 2018:557(7703):43–49. 10.1038/s41586-018-0063-9

Wase N, Abshire N, and Obata T. High-Throughput Profiling of Metabolic Phenotypes Using High-Resolution GC-MS. In. High-Throughput Plant Phenotyping, A Lorence and K Medina Jimenez, eds, Methods in Molecular Biology. (Springer US: New York, NY), pp. 235–260. 10.1007/978-1-0716-2537-8_19

Wen W, Li D, Li X, Gao Y, Li W, Li H, Liu J, Liu H, Chen W, Luo J, et al. Metabolome-based genome-wide association study of maize kernel leads to novel biochemical insights. Nat Commun. 2014:5(1):3438. 10.1038/ncomms4438

Wen W, Li K, Alseek S, Omranian N, Zhao L, Zhou Y, Xiao Y, Jin M, Yang N, Liu H, et al. Genetic Determinants of the Network of Primary Metabolism and Their Relationships to Plant Performance in a Maize Recombinant Inbred Line Population. Plant Cell. 2015:27(7):1839–56. 10.1105/tpc.15.00208

Yokoyama R, De Oliveira MVV, Kleven B, and Maeda HA. The entry reaction of the plant shikimate pathway is subjected to highly complex metabolite-mediated regulation. Plant Cell. 2021:33(3):671–696. 10.1093/plcell/koaa042

Yu J, Pressoir G, Briggs WH, Vroh Bi I, Yamasaki M, Doebley JF, McMullen MD, Gaut BS, Nielsen DM, Holland JB, et al. A unified mixed-model method for association mapping that accounts for multiple levels of relatedness. Nat Genet. 2006:38(2):203–208. 10.1038/ng1702

Yuan H, Cheung CYM, Hilbers PAJ, and Van Riel NAW. Flux Balance Analysis of Plant Metabolism: The Effect of Biomass Composition and Model Structure on Model Predictions. Front Plant Sci. 2016:7. 10.3389/fpls.2016.00537

Zhang X, Zhang S, Hao F, Lai X, Yu H, Huang Y, and Wang H. Expression, Purification and Properties of Shikimate Dehydrogenase from Mycobacterium Tuberculosis. BMB Rep. 2005:38(5):624–631. 10.5483/BMBRep.2005.38.5.624

Zheng L-Y, Guo X-S, He B, Sun L-J, Peng Y, Dong S-S, Liu T-F, Jiang S, Ramachandran S, Liu C-M, et al. Genome-wide patterns of genetic variation in sweet and grain sorghum (Sorghum bicolor). Genome Biol. 2011:12(11):R114. 10.1186/gb-2011-12-11-r114

Zhou S, Kremling KA, Bandillo N, Richter A, Zhang YK, Ahern KR, Artyukhin AB, Hui JX, Younkin GC, Schroeder FC, et al. Metabolome-Scale Genome-Wide Association Studies Reveal Chemical Diversity and Genetic Control of Maize Specialized Metabolites. Plant Cell. 2019:31(5):937–955. 10.1105/tpc.18.00772

Zhou X and Stephens M. Genome-wide efficient mixed-model analysis for association studies. Nat Genet. 2012:44(7):821–824. 10.1038/ng.2310

Zhou Z, Jiang Y, Wang Z, Gou Z, Lyu J, Li W, Yu Y, Shu L, Zhao Y, Ma Y, et al. Resequencing 302 wild and cultivated accessions identifies genes related to domestication and improvement in soybean. Nat Biotechnol. 2015:33(4):408– 414. 10.1038/nbt.3096

