## Supplementary doc 1 for "Natural Variation in Maize Shikimate Dehydrogenase Alters Enzyme Activity and Kernel Homoserine Accumulation"

**Supplementary doc 1:** pET21b with *Sad1*_Hap1 and pET21b with *Sad1*_Hap2 sequence

**Vector construct 1: DNA sequence of pET21b with Sad1_Hap1**

tggcgaatgggacgcgccctgtagcggcgcattaagcgcggcgggtgtggtggttacgcgcagcgtgaccgctacacttgccagcgccctagcgcccgctcctttcgctttcttcccttcctttctcgccacgttcgccggctttccccgtcaagctctaaatcgggggctccctttagggttccgatttagtgctttacggcacctcgaccccaaaaaacttgattagggtgatggttcacgtagtgggccatcgccctgatagacggtttttcgccctttgacgttggagtccacgttctttaatagtggactcttgttccaaactggaacaacactcaaccctatctcggtctattcttttgatttataagggattttgccgatttcggcctattggttaaaaaatgagctgatttaacaaaaatttaacgcgaattttaacaaaatattaacgtttacaatttcaggtggcacttttcggggaaatgtgcgcggaacccctatttgtttatttttctaaatacattcaaatatgtatccgctcatgagacaataaccctgataaatgcttcaataatattgaaaaaggaagagtatgagtattcaacatttccgtgtcgcccttattcccttttttgcggcattttgccttcctgtttttgctcacccagaaacgctggtgaaagtaaaagatgctgaagatcagttgggtgcacgagtgggttacatcgaactggatctcaacagcggtaagatccttgagagttttcgccccgaagaacgttttccaatgatgagcacttttaaagttctgctatgtggcgcggtattatcccgtattgacgccgggcaagagcaactcggtcgccgcatacactattctcagaatgacttggttgagtactcaccagtcacagaaaagcatcttacggatggcatgacagtaagagaattatgcagtgctgccataaccatgagtgataacactgcggccaacttacttctgacaacgatcggaggaccgaaggagctaaccgcttttttgcacaacatgggggatcatgtaactcgccttgatcgttgggaaccggagctgaatgaagccataccaaacgacgagcgtgacaccacgatgcctgcagcaatggcaacaacgttgcgcaaactattaactggcgaactacttactctagcttcccggcaacaattaatagactggatggaggcggataaagttgcaggaccacttctgcgctcggcccttccggctggctggtttattgctgataaatctggagccggtgagcgtgggtctcgcggtatcattgcagcactggggccagatggtaagccctcccgtatcgtagttatctacacgacggggagtcaggcaactatggatgaacgaaatagacagatcgctgagataggtgcctcactgattaagcattggtaactgtcagaccaagtttactcatatatactttagattgatttaaaacttcatttttaatttaaaaggatctaggtgaagatcctttttgataatctcatgaccaaaatcccttaacgtgagttttcgttccactgagcgtcagaccccgtagaaaagatcaaaggatcttcttgagatcctttttttctgcgcgtaatctgctgcttgcaaacaaaaaaaccaccgctaccagcggtggtttgtttgccggatcaagagctaccaactctttttccgaaggtaactggcttcagcagagcgcagataccaaatactgtccttctagtgtagccgtagttaggccaccacttcaagaactctgtagcaccgcctacatacctcgctctgctaatcctgttaccagtggctgctgccagtggcgataagtcgtgtcttaccgggttggactcaagacgatagttaccggataaggcgcagcggtcgggctgaacggggggttcgtgcacacagcccagcttggagcgaacgacctacaccgaactgagatacctacagcgtgagctatgagaaagcgccacgcttcccgaagggagaaaggcggacaggtatccggtaagcggcagggtcggaacaggagagcgcacgagggagcttccagggggaaacgcctggtatctttatagtcctgtcgggtttcgccacctctgacttgagcgtcgatttttgtgatgctcgtcaggggggcggagcctatggaaaaacgccagcaacgcggcctttttacggttcctggccttttgctggccttttgctcacatgttctttcctgcgttatcccctgattctgtggataaccgtattaccgcctttgagtgagctgataccgctcgccgcagccgaacgaccgagcgcagcgagtcagtgagcgaggaagcggaagagcgcctgatgcggtattttctccttacgcatctgtgcggtatttcacaccgcatatatggtgcactctcagtacaatctgctctgatgccgcatagttaagccagtatacactccgctatcgctacgtgactgggtcatggctgcgccccgacacccgccaacacccgctgacgcgccctgacgggcttgtctgctcccggcatccgcttacagacaagctgtgaccgtctccgggagctgcatgtgtcagaggttttcaccgtcatcaccgaaacgcgcgaggcagctgcggtaaagctcatcagcgtggtcgtgaagcgattcacagatgtctgcctgttcatccgcgtccagctcgttgagtttctccagaagcgttaatgtctggcttctgataaagcgggccatgttaagggcggttttttcctgtttggtcactgatgcctccgtgtaagggggatttctgttcatgggggtaatgataccgatgaaacgagagaggatgctcacgatacgggttactgatgatgaacatgcccggttactggaacgttgtgagggtaaacaactggcggtatggatgcggcgggaccagagaaaaatcactcagggtcaatgccagcgcttcgttaatacagatgtaggtgttccacagggtagccagcagcatcctgcgatgcagatccggaacataatggtgcagggcgctgacttccgcgtttccagactttacgaaacacggaaaccgaagaccattcatgttgttgctcaggtcgcagacgttttgcagcagcagtcgcttcacgttcgctcgcgtatcggtgattcattctgctaaccagtaaggcaaccccgccagcctagccgggtcctcaacgacaggagcacgatcatgcgcacccgtggggccgccatgccggcgataatggcctgcttctcgccgaaacgtttggtggcgggaccagtgacgaaggcttgagcgagggcgtgcaagattccgaataccgcaagcgacaggccgatcatcgtcgcgctccagcgaaagcggtcctcgccgaaaatgacccagagcgctgccggcacctgtcctacgagttgcatgataaagaagacagtcataagtgcggcgacgatagtcatgccccgcgcccaccggaaggagctgactgggttgaaggctctcaagggcatcggtcgagatcccggtgcctaatgagtgagctaacttacattaattgcgttgcgctcactgcccgctttccagtcgggaaacctgtcgtgccagctgcattaatgaatcggccaacgcgcggggagaggcggtttgcgtattgggcgccagggtggtttttcttttcaccagtgagacgggcaacagctgattgcccttcaccgcctggccctgagagagttgcagcaagcggtccacgctggtttgccccagcaggcgaaaatcctgtttgatggtggttaacggcgggatataacatgagctgtcttcggtatcgtcgtatcccactaccgagatatccgcaccaacgcgcagcccggactcggtaatggcgcgcattgcgcccagcgccatctgatcgttggcaaccagcatcgcagtgggaacgatgccctcattcagcatttgcatggtttgttgaaaaccggacatggcactccagtcgccttcccgttccgctatcggctgaatttgattgcgagtgagatatttatgccagccagccagacgcagacgcgccgagacagaacttaatgggcccgctaacagcgcgatttgctggtgacccaatgcgaccagatgctccacgcccagtcgcgtaccgtcttcatgggagaaaataatactgttgatgggtgtctggtcagagacatcaagaaataacgccggaacattagtgcaggcagcttccacagcaatggcatcctggtcatccagcggatagttaatgatcagcccactgacgcgttgcgcgagaagattgtgcaccgccgctttacaggcttcgacgccgcttcgttctaccatcgacaccaccacgctggcacccagttgatcggcgcgagatttaatcgccgcgacaatttgcgacggcgcgtgcagggccagactggaggtggcaacgccaatcagcaacgactgtttgcccgccagttgttgtgccacgcggttgggaatgtaattcagctccgccatcgccgcttccactttttcccgcgttttcgcagaaacgtggctggcctggttcaccacgcgggaaacggtctgataagagacaccggcatactctgcgacatcgtataacgttactggtttcacattcaccaccctgaattgactctcttccgggcgctatcatgccataccgcgaaaggttttgcgccattcgatggtgtccgggatctcgacgctctcccttatgcgactcctgcattaggaagcagcccagtagtaggttgaggccgttgagcaccgccgccgcaaggaatggtgcatgcaaggagatggcgcccaacagtcccccggccacggggcctgccaccatacccacgccgaaacaagcgctcatgagcccgaagtggcgagcccgatcttccccatcggtgatgtcggcgatataggcgccagcaaccgcacctgtggcgccggtgatgccggccacgatgcgtccggcgtagaggatcgagatctcgatcccgcgaaattaatacgactcactataggggaattgtgagcggataacaattcccctctagaaataattttgtttaactttaagaaggagatatacatatggctagcatggtctgcgtgccggcgacggcgcgggcgccgcgggagatggcggcggagctggccgccgccgccgcgctcggcgctgacgtggcggagcttcgcctcgactgcctcgccgggttcgcgccgcgcagggacctgcccgtcatcctcgccaagccccgcccgctccccgccctcgtcacctacaggcccaagtgggaaggaggcgaatacgaaggcgatgacgaatcacggtttgaggctctgctattggcaatggagctgggagctgaatatgtggatgtcgagcttaaggtggctgacaaatttatgaaacttatttctgggaggaaacctgataactgtaaacttatagtttcatcccacaactatgagaccactccatcgtccgaggaacttgcaaatttggtggctcagattcaagcaacgggggctgatatcgtgaaaatagctacaaccgctactgaaattgttgatgtggcaaaaatgtttcaaatacttgttcactgccaggaaaagcaggtgccaatcattgggctggtgatgaacgacagaggttttatttctcgggttctttgcccaaaatatggtggattccttacttttgggtcactcaagaaaggaaaagagtctgcacctgcacagccaactgctgcagacttgataaatctgtacaatattaggcagatagggccagacactaaggtctttggtataattggtaaaccagttggccatagcaaaagcccaattttgcataatgaagctttcagatctgtgggtttcaacgctgtgtatgtgccatttttggtggatgacttggctaaatttcttgatacatactcttcaccagactttgctggcttcagttgcacaattccacacaaagaagctgctgttaggtgctgtgacgaggtcgatcccgttgccagggacattggagctgttaacacaattgttagaagacctgatggaaagcttgttggctataatactgactatgttggtgctatatctgctattgaggatggaataaaagcatcagaatcagaaccaacagatccagacaaatcaccactggctggaaggctttttgttgttataggggctggtggtgcgggaaaagcactagcatatggggcaaaagagaaaggagcaagagttgtaattgcaaaccgtacctttgcacgagcacaagaacttgccaacttaattggtgggcctgcattgactcttgcggatttggaaaactaccatccagaggaagggatgattcttgcaaatacaacagccattggaatgcatccaaatgtgaatgaaactcctctatctaagcaagcactcagatcttatgctgttgtgtttgatgcggtctacacaccaaaagagaccagacttctccgagaagctgcagaatgtggagccacagttgttagtggtctggagatgtttatacgccaagctatgggccagtttgagcatttcacgggcatgccagctccagatagcttgatgcgtgatattgttctgacaaagacatacaaacaatacaccatttcaacaatgtttaagagttcttactcctctcatcaaggctgcaaggatttggacgattacaaattccgagcaattgcaagttcattggacaatcctaacaccatgaagcctcgtctgGGCGGCGGCTCCgagaacctatatttccagggatacccatacgacgtaccagattacgctctcgagcaccaccaccaccaccactgagatccggctgctaacaaagcccgaaaggaagctgagttggctgctgccaccgctgagcaataactagcataaccccttggggcctctaaacgggtcttgaggggttttttgctgaaaggaggaactatatccggat

**Vector construct 2: DNA sequence of pET21b with Sad1_Hap2**

TGGCGAATGGGACGCGCCCTGTAGCGGCGCATTAAGCGCGGCGGGTGTGGTGGTTACGCGCAGCGTGACCGCTACACTTGCCAGCGCCCTAGCGCCCGCTCCTTTCGCTTTCTTCCCTTCCTTTCTCGCCACGTTCGCCGGCTTTCCCCGTCAAGCTCTAAATCGGGGGCTCCCTTTAGGGTTCCGATTTAGTGCTTTACGGCACCTCGACCCCAAAAAACTTGATTAGGGTGATGGTTCACGTAGTGGGCCATCGCCCTGATAGACGGTTTTTCGCCCTTTGACGTTGGAGTCCACGTTCTTTAATAGTGGACTCTTGTTCCAAACTGGAACAACACTCAACCCTATCTCGGTCTATTCTTTTGATTTATAAGGGATTTTGCCGATTTCGGCCTATTGGTTAAAAAATGAGCTGATTTAACAAAAATTTAACGCGAATTTTAACAAAATATTAACGTTTACAATTTCAGGTGGCACTTTTCGGGGAAATGTGCGCGGAACCCCTATTTGTTTATTTTTCTAAATACATTCAAATATGTATCCGCTCATGAGACAATAACCCTGATAAATGCTTCAATAATATTGAAAAAGGAAGAGTATGAGTATTCAACATTTCCGTGTCGCCCTTATTCCCTTTTTTGCGGCATTTTGCCTTCCTGTTTTTGCTCACCCAGAAACGCTGGTGAAAGTAAAAGATGCTGAAGATCAGTTGGGTGCACGAGTGGGTTACATCGAACTGGATCTCAACAGCGGTAAGATCCTTGAGAGTTTTCGCCCCGAAGAACGTTTTCCAATGATGAGCACTTTTAAAGTTCTGCTATGTGGCGCGGTATTATCCCGTATTGACGCCGGGCAAGAGCAACTCGGTCGCCGCATACACTATTCTCAGAATGACTTGGTTGAGTACTCACCAGTCACAGAAAAGCATCTTACGGATGGCATGACAGTAAGAGAATTATGCAGTGCTGCCATAACCATGAGTGATAACACTGCGGCCAACTTACTTCTGACAACGATCGGAGGACCGAAGGAGCTAACCGCTTTTTTGCACAACATGGGGGATCATGTAACTCGCCTTGATCGTTGGGAACCGGAGCTGAATGAAGCCATACCAAACGACGAGCGTGACACCACGATGCCTGCAGCAATGGCAACAACGTTGCGCAAACTATTAACTGGCGAACTACTTACTCTAGCTTCCCGGCAACAATTAATAGACTGGATGGAGGCGGATAAAGTTGCAGGACCACTTCTGCGCTCGGCCCTTCCGGCTGGCTGGTTTATTGCTGATAAATCTGGAGCCGGTGAGCGTGGGTCTCGCGGTATCATTGCAGCACTGGGGCCAGATGGTAAGCCCTCCCGTATCGTAGTTATCTACACGACGGGGAGTCAGGCAACTATGGATGAACGAAATAGACAGATCGCTGAGATAGGTGCCTCACTGATTAAGCATTGGTAACTGTCAGACCAAGTTTACTCATATATACTTTAGATTGATTTAAAACTTCATTTTTAATTTAAAAGGATCTAGGTGAAGATCCTTTTTGATAATCTCATGACCAAAATCCCTTAACGTGAGTTTTCGTTCCACTGAGCGTCAGACCCCGTAGAAAAGATCAAAGGATCTTCTTGAGATCCTTTTTTTCTGCGCGTAATCTGCTGCTTGCAAACAAAAAAACCACCGCTACCAGCGGTGGTTTGTTTGCCGGATCAAGAGCTACCAACTCTTTTTCCGAAGGTAACTGGCTTCAGCAGAGCGCAGATACCAAATACTGTCCTTCTAGTGTAGCCGTAGTTAGGCCACCACTTCAAGAACTCTGTAGCACCGCCTACATACCTCGCTCTGCTAATCCTGTTACCAGTGGCTGCTGCCAGTGGCGATAAGTCGTGTCTTACCGGGTTGGACTCAAGACGATAGTTACCGGATAAGGCGCAGCGGTCGGGCTGAACGGGGGGTTCGTGCACACAGCCCAGCTTGGAGCGAACGACCTACACCGAACTGAGATACCTACAGCGTGAGCTATGAGAAAGCGCCACGCTTCCCGAAGGGAGAAAGGCGGACAGGTATCCGGTAAGCGGCAGGGTCGGAACAGGAGAGCGCACGAGGGAGCTTCCAGGGGGAAACGCCTGGTATCTTTATAGTCCTGTCGGGTTTCGCCACCTCTGACTTGAGCGTCGATTTTTGTGATGCTCGTCAGGGGGGCGGAGCCTATGGAAAAACGCCAGCAACGCGGCCTTTTTACGGTTCCTGGCCTTTTGCTGGCCTTTTGCTCACATGTTCTTTCCTGCGTTATCCCCTGATTCTGTGGATAACCGTATTACCGCCTTTGAGTGAGCTGATACCGCTCGCCGCAGCCGAACGACCGAGCGCAGCGAGTCAGTGAGCGAGGAAGCGGAAGAGCGCCTGATGCGGTATTTTCTCCTTACGCATCTGTGCGGTATTTCACACCGCATATATGGTGCACTCTCAGTACAATCTGCTCTGATGCCGCATAGTTAAGCCAGTATACACTCCGCTATCGCTACGTGACTGGGTCATGGCTGCGCCCCGACACCCGCCAACACCCGCTGACGCGCCCTGACGGGCTTGTCTGCTCCCGGCATCCGCTTACAGACAAGCTGTGACCGTCTCCGGGAGCTGCATGTGTCAGAGGTTTTCACCGTCATCACCGAAACGCGCGAGGCAGCTGCGGTAAAGCTCATCAGCGTGGTCGTGAAGCGATTCACAGATGTCTGCCTGTTCATCCGCGTCCAGCTCGTTGAGTTTCTCCAGAAGCGTTAATGTCTGGCTTCTGATAAAGCGGGCCATGTTAAGGGCGGTTTTTTCCTGTTTGGTCACTGATGCCTCCGTGTAAGGGGGATTTCTGTTCATGGGGGTAATGATACCGATGAAACGAGAGAGGATGCTCACGATACGGGTTACTGATGATGAACATGCCCGGTTACTGGAACGTTGTGAGGGTAAACAACTGGCGGTATGGATGCGGCGGGACCAGAGAAAAATCACTCAGGGTCAATGCCAGCGCTTCGTTAATACAGATGTAGGTGTTCCACAGGGTAGCCAGCAGCATCCTGCGATGCAGATCCGGAACATAATGGTGCAGGGCGCTGACTTCCGCGTTTCCAGACTTTACGAAACACGGAAACCGAAGACCATTCATGTTGTTGCTCAGGTCGCAGACGTTTTGCAGCAGCAGTCGCTTCACGTTCGCTCGCGTATCGGTGATTCATTCTGCTAACCAGTAAGGCAACCCCGCCAGCCTAGCCGGGTCCTCAACGACAGGAGCACGATCATGCGCACCCGTGGGGCCGCCATGCCGGCGATAATGGCCTGCTTCTCGCCGAAACGTTTGGTGGCGGGACCAGTGACGAAGGCTTGAGCGAGGGCGTGCAAGATTCCGAATACCGCAAGCGACAGGCCGATCATCGTCGCGCTCCAGCGAAAGCGGTCCTCGCCGAAAATGACCCAGAGCGCTGCCGGCACCTGTCCTACGAGTTGCATGATAAAGAAGACAGTCATAAGTGCGGCGACGATAGTCATGCCCCGCGCCCACCGGAAGGAGCTGACTGGGTTGAAGGCTCTCAAGGGCATCGGTCGAGATCCCGGTGCCTAATGAGTGAGCTAACTTACATTAATTGCGTTGCGCTCACTGCCCGCTTTCCAGTCGGGAAACCTGTCGTGCCAGCTGCATTAATGAATCGGCCAACGCGCGGGGAGAGGCGGTTTGCGTATTGGGCGCCAGGGTGGTTTTTCTTTTCACCAGTGAGACGGGCAACAGCTGATTGCCCTTCACCGCCTGGCCCTGAGAGAGTTGCAGCAAGCGGTCCACGCTGGTTTGCCCCAGCAGGCGAAAATCCTGTTTGATGGTGGTTAACGGCGGGATATAACATGAGCTGTCTTCGGTATCGTCGTATCCCACTACCGAGATATCCGCACCAACGCGCAGCCCGGACTCGGTAATGGCGCGCATTGCGCCCAGCGCCATCTGATCGTTGGCAACCAGCATCGCAGTGGGAACGATGCCCTCATTCAGCATTTGCATGGTTTGTTGAAAACCGGACATGGCACTCCAGTCGCCTTCCCGTTCCGCTATCGGCTGAATTTGATTGCGAGTGAGATATTTATGCCAGCCAGCCAGACGCAGACGCGCCGAGACAGAACTTAATGGGCCCGCTAACAGCGCGATTTGCTGGTGACCCAATGCGACCAGATGCTCCACGCCCAGTCGCGTACCGTCTTCATGGGAGAAAATAATACTGTTGATGGGTGTCTGGTCAGAGACATCAAGAAATAACGCCGGAACATTAGTGCAGGCAGCTTCCACAGCAATGGCATCCTGGTCATCCAGCGGATAGTTAATGATCAGCCCACTGACGCGTTGCGCGAGAAGATTGTGCACCGCCGCTTTACAGGCTTCGACGCCGCTTCGTTCTACCATCGACACCACCACGCTGGCACCCAGTTGATCGGCGCGAGATTTAATCGCCGCGACAATTTGCGACGGCGCGTGCAGGGCCAGACTGGAGGTGGCAACGCCAATCAGCAACGACTGTTTGCCCGCCAGTTGTTGTGCCACGCGGTTGGGAATGTAATTCAGCTCCGCCATCGCCGCTTCCACTTTTTCCCGCGTTTTCGCAGAAACGTGGCTGGCCTGGTTCACCACGCGGGAAACGGTCTGATAAGAGACACCGGCATACTCTGCGACATCGTATAACGTTACTGGTTTCACATTCACCACCCTGAATTGACTCTCTTCCGGGCGCTATCATGCCATACCGCGAAAGGTTTTGCGCCATTCGATGGTGTCCGGGATCTCGACGCTCTCCCTTATGCGACTCCTGCATTAGGAAGCAGCCCAGTAGTAGGTTGAGGCCGTTGAGCACCGCCGCCGCAAGGAATGGTGCATGCAAGGAGATGGCGCCCAACAGTCCCCCGGCCACGGGGCCTGCCACCATACCCACGCCGAAACAAGCGCTCATGAGCCCGAAGTGGCGAGCCCGATCTTCCCCATCGGTGATGTCGGCGATATAGGCGCCAGCAACCGCACCTGTGGCGCCGGTGATGCCGGCCACGATGCGTCCGGCGTAGAGGATCGAGATCTCGATCCCGCGAAATTAATACGACTCACTATAGGGGAATTGTGAGCGGATAACAATTCCCCTCTAGAAATAATTTTGTTTAACTTTAAGAAGGAGATATACATATGGCTAGCATGGTCTGCGTGCCGGCGACGGCGCGGGCGCCGCGGGAGATGGCGGCGGAGCTGGCCGCCGCCGCCGCGtTCGGCGCTGACGTGGCGGAGCTTCGCCTCGACTGCCTCGCCGGGTTCGCGCCGCGCAGGGACCTGCCCGTCATCCTCGCCAAGCCCCGCCCGCTCCCCGCCCTCGTCACCTACAGGCCCAAGTGGGAAGGAGGCGAATACGAAGGCGATGACGAATCACGGTTTGAGGCTCTGCTATTGGCAATGGAGCTGGGAGCTGAATATGTGGATGTCGAGCTTAAGGTGGCTGACAAATTTATGAAACTTATTTCTGGGAGGAAACCTGATAACTGTAAACTTATAGTTTCATCCCACAACTATGAGACCACTCCATCGTCCGAGGAACTTGCAAATTTGGTGGCTCAGATTCAAGCAACGGGGGCTGATATCGTGAAAATAGCTACAACCGCTACTGAAATTGTTGATGTGGCAAAAATGTTTCAAATACTTGTTCACTGCCAGGAAAAGCAGGTGCCAATCATTGGGCTGGTGATGAACGACAGAGGTTTTATTTCTCGGGTTCTTTGCCCAAAATATGGTGGATTCCTTACTTTTGGGTCACTCAAGAAAGGAAAAGAGTCTGCACCTGCACAGCCAACTGCTGCAGACTTGATAAATCTGTACAATATTAGGCAGATAGGGCCAGACACTAAGGTCTTTGGTATAATTGGTAAACCAGTTGGCCATAGCAAAAGCCCAATTTTGCATAATGAAGCTTTCAGATCTGTGGGTTTCAACGCTGTGTATGTGCCATTTTTGGTGGATGACTTGGCTAAATTTCTTGATACATACTCTTCACCAGACTTTGCTGGCTTCAGTTGCACAATTCCACACAAAGAAGCTGCTGTTAGGTGCTGTGACGAGGTCGATCCCGTTGCCAGGGACATTGGAGCTGTTAACACAATTGTTAGAAGACCTGATGGAAAGCTTGTTGGCTATAATACTGACTATGTTGGTGCTATATCTGCTATTGAGGATGGAATAAAAGCATCAGAATCAGAACCAACAGATCCAGACAAATCACCACTGGCTGGAAGGCTTTTTGTTGTTATAGGGGCTGGTGGTGCGGGAAAAGCACTAGCATATGGGGCAAAAGAGAAAGGAGCAAGAGTTGTAATTGCAAACCGTACCTTTGCACGAGCACAAGAACTTGCCAACTTAATTGGTGGGCCTGCATTGACTCTTGCGGATTTGGAAAACTACCATCCAGAGGAAGGGATGATTCTTGCAAATACAACAGCCATTGGAATGCATCCAAATGTGAATGAAACTCCTCTATCTAAGCAAGCACTCAGATCTTATGCTGTTGTGTTTGATGCGGTCTACACACCAAAAGAGACCAGACTTCTCCGAGAAGCTGCAGAATGTGGAGCCACAGTTGTTAGTGGTCTGGAGATGTTTATACGCCAAGCTATGGGCCAGTTTGAGCATTTCACGGGCATGCCAGCTtCAGATAGCTTGATGCGTGATATTGTTCTGACAAAGACATACAAACtATACACCATTTCAACAATGTTTAAGAGTTCTTACTCCTCTCATCAAGGCTGCAAGGATTTGGACGATTACAAATTCCGAGCAATTGCAAGTTCATTGGACAATCCTAACACCATGAAGCCTCGTCTGGGCGGCGGCTCCgagaacctatatttccagggaTACCCATACGACGTACCAGATTACGCTCTCGAGCACCACCACCACCACCACTGAGATCCGGCTGCTAACAAAGCCCGAAAGGAAGCTGAGTTGGCTGCTGCCACCGCTGAGCAATAACTAGCATAACCCCTTGGGGCCTCTAAACGGGTCTTGAGGGGTTTTTTGCTGAAAGGAGGAACTATATCCGGAT

Note: Yellow highlighted DNA sequences are Sad1_Hap1 and Sad1_Hap2 respectively in Vector construct 1 and Vector construct 2. Red highlighted nucleotide is the changes in DNA between Sad1_Hap1 and Sad1_Hap2.
